# Blocked transcription-translation complexes are rescued by transcript release followed by *trans*-translation

**DOI:** 10.64898/2026.05.06.723219

**Authors:** Stephanie L. Leedom, Kenneth C. Keiler

**Affiliations:** Department of Molecular Biosciences, The University of Texas at Austin, Austin, Texas, United States of America; LaMontagne Center for Infectious Diseases, The University of Texas at Austin, Austin, Texas, United States of America

## Abstract

Bacterial ribosomes initiate translation while the nascent transcript is still engaged with RNA polymerase, creating a risk that translating ribosomes will become trapped if RNA polymerase is blocked before transcription of the stop codon. Here we show that DNA-binding proteins block RNA polymerase and translating ribosomes *in vitro* and *in vivo*. Translating ribosomes are rescued after they trigger release of the nascent transcript from RNA polymerase. Following release of the transcript, ribosomes translate to the 3’ end of the mRNA and are rescued by *trans*-translation. This mechanism allows rescue of all components of blocked transcription-translation reactions.

## Introduction

The bacterial RNA polymerase elongation complex is highly stable, allowing long mRNA transcripts to be efficiently synthesized (1,2). However, when an elongating RNA polymerase is blocked, for example by a DNA-binding protein or a DNA lesion, there must be a mechanism to release the transcript so the RNA polymerase can be recycled. Because trailing RNA polymerases do not remove blocked RNA polymerases (3), many RNA polymerases transcribing the same gene could accumulate behind a single blockage. Potential problems for the cell are compounded by translation of the nascent mRNA, because ribosomes would also stall behind a blocked polymerase. Studies of transcription and translation of *Escherichia coli lacZ* indicate that each gene can quickly accumulate >30 RNA polymerases and >1200 ribosomes (4), so blocked RNA polymerases could create a strain on cellular resources.

The fate of blocked RNA polymerase-ribosome complexes is not clear. In one *in vitro* study, when RNA polymerase was blocked by a pyrimidine dimer and a translating ribosome was stepped forward by serial addition of cognate tRNAs, the elongation complex was destroyed (3). This result demonstrated that a translating ribosome can disrupt some blocked RNA polymerases. However, these experiments could not determine whether the RNA polymerase was removed from the DNA by the translating ribosome or if the nascent transcript was released from RNA polymerase. In addition, ribosomes terminated translation during the experiment because a stop codon was present on the nascent transcript, so the fate of ribosomes behind an RNA polymerase blocked within an open reading frame has not been investigated. Here, we sought to determine the fate of ribosomes in blocked transcription-translation complexes in living cells and gain insight into the mechanism for rescuing these blocked complexes.

Several mechanisms for rescuing ribosomes in bacteria are known. Ribosomes that translate to the 3’ terminus of an mRNA without terminating are typically rescued by *trans*-translation (5). During *trans*-translation, transfer-messenger RNA (tmRNA) is used first like a tRNA and then as a template for translation to add a proteolysis signal, the SsrA tag, to the nascent polypeptide (6,7). The ribosome terminates at a stop codon within tmRNA and is recycled (6). *trans*-Translation also promotes degradation of the mRNA that the ribosome stalled on, so the nascent polypeptide, the ribosome, and the mRNA are all removed (6,8,9). Some bacteria also have proteins that can act as alternative ribosome rescue factors to release ribosomes stalled at the 3’ end of an mRNA (10–15). For example, *E. coli* ArfA recruits RF2 to hydrolyze peptidyl-tRNA on ribosomes stalled at the 3’ end of an mRNA, releasing the ribosome and the untagged nascent polypeptide (16). *trans*-Translation and the alternative ribosome rescue factors recognize the absence of mRNA at the leading edge of the ribosome and are highly inefficient on ribosomes stalled in the middle of a transcript (17). Ribosomes stalled in the middle of a transcript can be targeted to *trans*-translation by exo- or endonucleases, including SrmB, a nuclease that cuts the mRNA between two collided ribosomes (18). Stalled ribosomes can also be split by factors such as MutS2 or HrpA to remove them from the mRNA and recycle them (19,20). No nucleases are known to specifically target collided ribosome-RNA polymerase complexes.

To discover how ribosomes trapped by a blocked RNA polymerase are rescued, we used dCas9 to generate sequence-specific transcription blocks *in vivo* and *in vitro*. Four possible fates for transcription-translation complexes blocked by a DNA-binding protein are shown in Figure 1. First, the transcription-translation complex might be stable *in vivo*, with little rescue. Second, cleavage of the mRNA by an endonuclease might target the ribosome to *trans*-translation, or tmRNA or one of the alternative ribosome rescue factors might be able to rescue these blocked ribosomes despite mRNA extending past the leading edge of the ribosome. Third, the ribosome might knock RNA polymerase off the DNA and translate to the 3’ end of the transcript before being rescued. Finally, the nascent transcript might be released from RNA polymerase by the colliding ribosome, followed by translation to the 3’ end of the mRNA and rescue by *trans*-translation.

**Fig 1.**
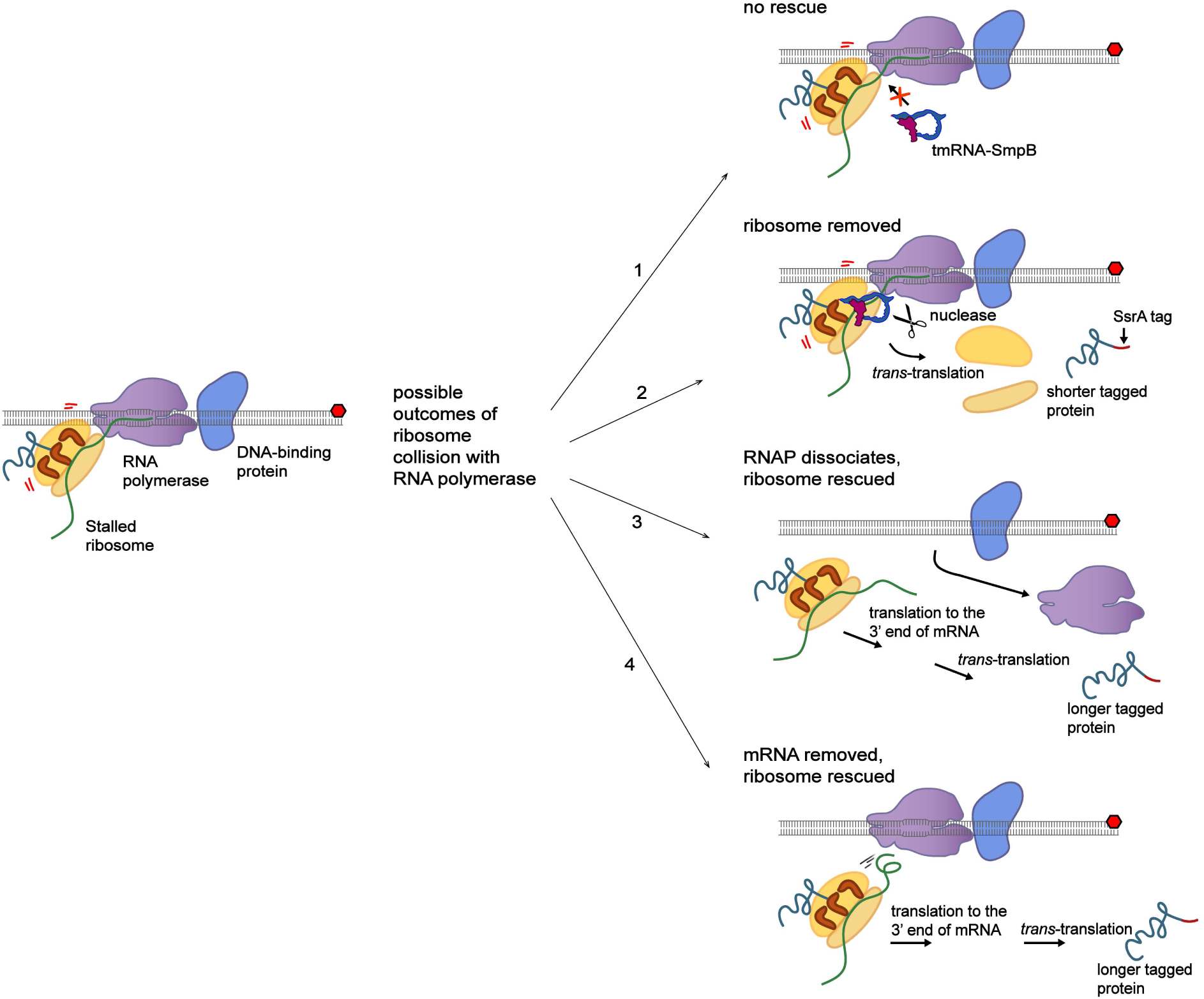
Possible outcomes of a blocked transcription-translation complex. A transcription-translation complex roadblocked by a DNA-binding protein is diagrammed on the left and four possible outcomes for the stalled machinery are shown. *Possibility 1*: There is no rescue and all components remain collided on the DNA and nascent transcript. *Possibility 2*: The ribosome is rescued after collision with RNA polymerase through cleavage of the mRNA by an unknown endonuclease or unusual activity by tmRNA-SmpB. *Possibility 3*: RNA polymerase dissociates from the DNA, releasing the transcript and allowing the ribosome to translate to the 3’ end, where it is rescued by *trans*-translation. *Possibility 4*: The ribosome pulls the mRNA out of RNA polymerase and translates to the 3’ end of the truncated transcript, where it is rescued by *trans*-translation. The tagged protein generated in *Possibility 2* is shorter than the tagged proteins generated in *Possibilities 3 and 4* because the ribosome stops translating where it collides with RNA polymerase instead of continuing to the 3’ end that is in the polymerase active site.

By examining the proteins that are produced from blocked transcription-translation complexes and the fate of the transcript and RNA polymerase, we find that the ribosome triggers release of the nascent transcript from the blocked RNA polymerase, followed by translation to the 3’ end of the mRNA and *trans*-translation (possibility 4 in Fig. 1). Transcription-translation blocks generated by the natural DNA-binding protein GalR or DNA-damaging agents appear to be rescued in the same way, indicating that this mechanism is broadly used to prevent accumulation of RNA polymerases and ribosomes behind stable transcription blocks.

## Results

### *trans*-Translation rescues ribosomes from blocked transcription-translation complexes *in vivo*

We used dCas9 as a high-affinity, site-specific DNA-binding protein to investigate the outcomes of blocked transcription-translation complexes. dCas9 is a catalytically inactive mutant of *Streptococcus pyogenes* Cas9 that maintains its ability to bind DNA at a sequence defined by the associated sgRNA, but does not cleave DNA due to mutations in its endonuclease domains (21). dCas9-sgRNA binds DNA with a dissociation constant (K_d_) in the low nanomolar range and blocks RNA polymerase efficiently (22–24). To determine if *trans*-translation rescues ribosomes when RNA polymerase is blocked *in vivo*, we designed a sgRNA to target a gene encoding FLAG-EGFP and assayed tagging using an *E. coli* strain in which the wild-type SsrA tag sequence (ANDENYALAA) is replaced with AND<u>HHHHHHD</u> (SsrA-H6D). The SsrA-H6D tag does not target proteins for rapid proteolysis, and proteins with the SsrA-H6D tag can be detected by western blotting for the H6 epitope (25). When FLAG-EGFP, dCas9, and sgRNA were expressed in the strain with SsrA-H6D, two truncated proteins were detected by western blotting with anti-FLAG antibody (Fig. 2A). The larger of the two truncated proteins was also detected on western blots with Penta-His antibody, indicating that it had the SsrA-H6D tag (Fig. 2B). Only the smaller truncated protein was observed in anti-FLAG blots when the experiment was performed in Δ*ssrA* cells, which do not have tmRNA and cannot perform *trans*-translation (Fig. 2A). Tagged truncated FLAG-EGFP was not observed when the PAM site in the FLAG-EGFP gene was mutated, confirming that the observed tagging is caused by dCas9 binding (Fig. S1). These results indicate that FLAG-EGFP is tagged by *trans*-translation *in vivo* when transcription is blocked by dCas9.

**Fig 2.**
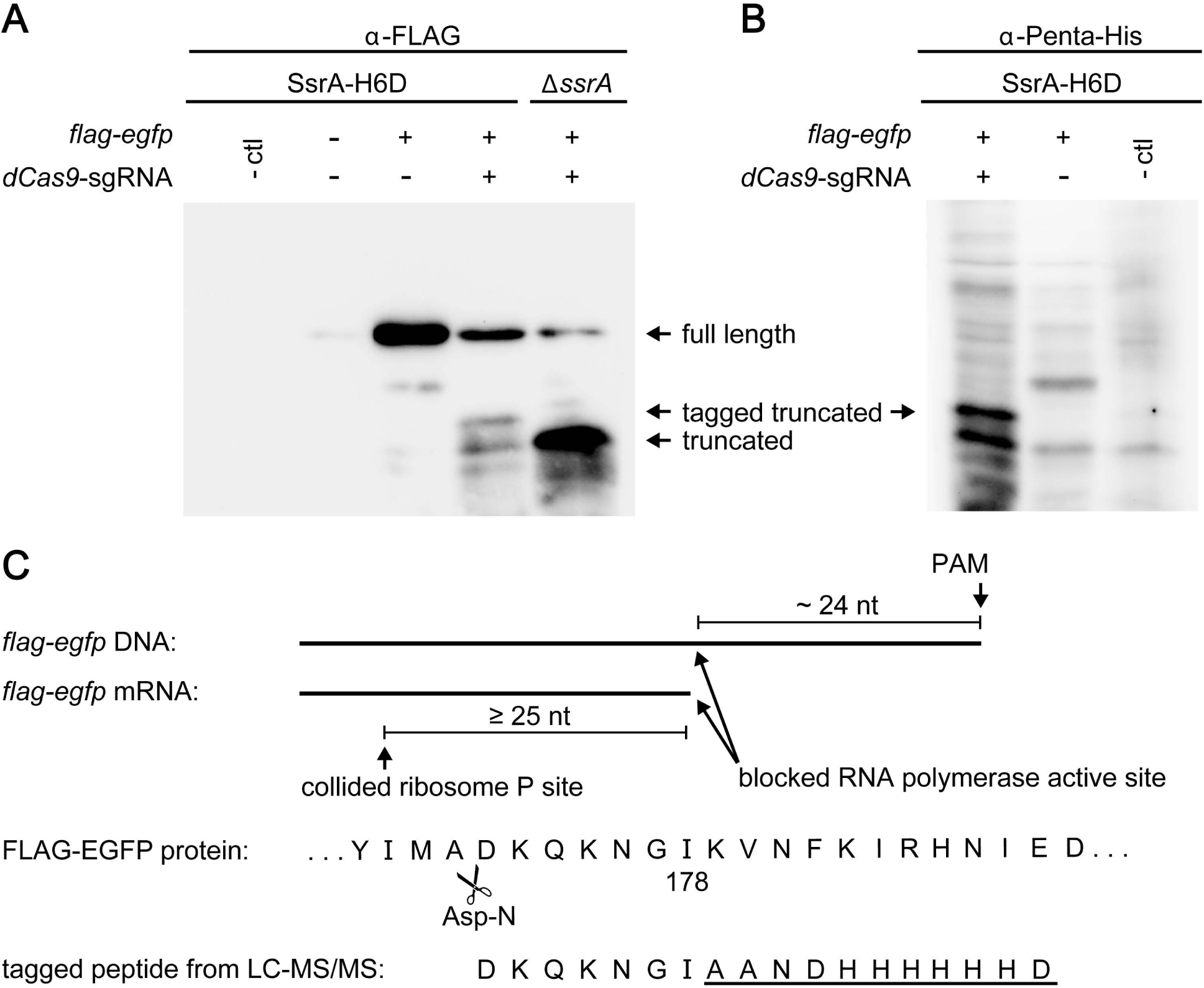
Blocking transcription with dCas9 triggers ribosome rescue by *trans*-translation. FLAG-EGFP and dCas9-sgRNA were expressed in *E. coli* containing a variant tmRNA with the SsrA-H6D tag (ANDHHHHHHD) or with no tmRNA (Δ*ssrA*) and proteins were analyzed by western blotting with (*A*) α-FLAG antibody to visualize all FLAG-EGFP or (*B*) by Penta-His antibody to visualize all SsrA-tagged proteins. Tables above the blots indicate which genes were induced. Strains with no plasmids are included as a negative control (-ctl). The mobilities of FLAG-EGFP bands are indicated. Representative results from three biological repeats are shown. (*C*) FLAG-tagged proteins from the SsrA-H6D strain expressing FLAG-EGFP and dCas9-sgRNA were purified, digested with Asp-N protease, and analyzed by LC-MS/MS. The SsrA-H6D tag was observed after I178 of FLAG-EGFP. The predicted relative locations of the PAM site, the RNA polymerase active site, and the ribosomal P site in the blocked complex are indicated in the diagram, and sequence of full-length FLAG-EGFP in this region is shown. The codon for I178 would be at or near the RNA polymerase active site. The SsrA-H6D tagged peptide from LC-MS/MS is shown below. The first alanine in the AANDHHHHHHD peptide sequence comes from the tRNA-like activity of tmRNA.

**Fig S1. dCas9-sgRNA truncation and tagging by tmRNA-SmpB requires a functional PAM site.** FLAG-EGFP with either the native PAM site (TGG) or the PAM site mutated to TTT was expressed with dCas9-sgRNA in *E. coli* SsrA-H6D and detected with α-FLAG antibody. Strains expressing truncated FLAG-EGFP (residues 1-179) and truncated FLAG-EGFP with the SsrA-H6D tag at its C-terminus are shown as size markers.

We identified where the SsrA-H6D tag was added to FLAG-EGFP *in vivo* by pulling down FLAG-tagged proteins, digesting them with Asp-N protease, and mapping the fragments using LC-MS/MS (Fig. 2C, & Supplementary Figs. S2 & S3). These data showed that both full-length FLAG-EGFP and FLAG-EGFP with the SsrAH6D tag after Ile178 were present, consistent with bands seen in the western blot. Ile178 is encoded 8 codons upstream of the PAM site (Fig. 2C, Table S1). Optical trapping studies of RNA polymerase collided with dCas9 show ∼24 nucleotides between the PAM site and the active site of RNA polymerase (26), and studies of ribosomes collided with RNA polymerase indicate that the distance between the RNA polymerase active site and the ribosomal P site is ≥25 nucleotides (27,28). Because the Ile178 tagging site is only 8 codons upstream of the PAM site, ribosomes are translating mRNA that extends to the RNA polymerase active site in the blocked complex before *trans*-translation occurs (Fig. 2C). These results indicate that the nascent transcript is released from RNA polymerase with the ribosome still engaged, allowing the ribosome to translate all the way to the 3’ end of the mRNA (Possibility 3 or 4 from Figure 1).

**Fig S2. Fragment ions of FLAG-EGFP.** The predicted fragment ions for the peptide sequence DKQKNGIKVNFKIRHNIE (diagrammed in Figure 2C) are shown. Colored boxes indicate ions that were recovered in the MS/MS spectrum.

**Fig S3. Fragment ions of tagged FLAG-EGFP.** The predicted fragment ions for the tagged peptide sequence DKQKNGIAANDHHHHHHD (diagrammed in Figure 2C) are shown. Colored boxes indicate ions that were recovered in the MS/MS spectrum.

**Table S1. List of peptides identified by LC-MS/MS.**

Shorter proteins containing both the FLAG epitope and the SsrA-H6D tag were not detected on western blots and there were no alternative tagging sites above the limit of detection in the LC-MS/MS dataset, indicating that nuclease activity on the mRNA between RNA polymerase and ribosome, in the A site of the ribosome, and between blocked ribosomes is rare prior to *trans*-translation (Fig. 2, Table S1). In cells deleted for *ssrA*, protein from blocked ribosomes is released without *trans*-translation (Figure 2). This protein likely derives from ArfA activity on ribosomes that have translated to the 3’ end of the truncated transcript. ArfA is regulated by *trans*-translation and is normally tagged and degraded when *trans*-translation is active (29). ArfA protein is more abundant when tmRNA-H6D is present because proteins tagged by SsrA-H6D are degraded more slowly than proteins tagged with the wild-type SsrA tag (25,30). In the Δ*ssrA* strain, ArfA is the main mechanism for rescuing stalled ribosomes. Unlike *trans*-translation, ArfA activity does not target the mRNA for degradation (10). The increased abundance of truncated FLAG-EGFP in the Δ*ssrA* strain in Figure 2A is likely due to a combination of efficient rescue of ribosomes at the 3’ end of the truncated mRNA by ArfA and slower degradation of the truncated transcripts, allowing more translation.

To test whether *trans*-translation also rescues ribosomes when RNA polymerase is blocked by DNA damage, we used western blotting to detect SsrA-H6D tagging in cells treated with sub-lethal concentrations of the DNA crosslinker mitomycin C (MMC). MMC alkylates CG sequences in DNA and interferes with transcription *in vivo* (31,32). Cells treated with MMC had 24 ± 11% more SsrA-H6D tagging than untreated cells, consistent with tagging after transcription-translation blocks at DNA damage sites (Fig. S4). Previous work has shown that mutants lacking *trans*-translation activity are hypersensitive to MMC (33), consistent with MMC generating *trans*-translation substrates as observed here.

**Fig S4. SsrA-H6D tagging in cells treated with MMC.** (*A*) *E. coli* containing the SsrA-H6D tag was treated with MMC and the percent tagging relative to untreated was determined by western blot with Penta-His antibody. Tagging in the untreated sample was defined as 100%. (*B*) Bar plots showing the percent tagging and total protein (determined by Ponceau staining) of treated cells relative to untreated. Data are shown as the mean ± standard deviation of three biological replicates. The total protein was determined by densitometry with the untreated sample set to 100%.

### Ribosomes behind a blocked RNA polymerase trigger release of the nascent transcript

To investigate how *trans*-translation rescues ribosomes from blocked transcription-translation complexes, we examined the reaction *in vitro* using purified components. When we incubated dCas9-sgRNA and DNA template before *in vitro* transcription with *E. coli* RNA polymerase, almost all the transcript was truncated to a size consistent with blocking RNA polymerase elongation just before the dCas9 binding site (Fig. 3A). Likewise, when we incubated dCas9-sgRNA with DNA template before *in vitro* transcription-translation, a truncated protein was produced (Fig. 3B). This result indicates that ribosomes could not push RNA polymerase past the dCas9 block to produce full-length mRNA and protein. Truncated protein with a lower gel mobility was detected when tmRNA-SmpB was included in the transcription-translation reaction, consistent with *trans*-translation adding the 1.1 kDa SsrA tag after translation of the truncated mRNA (Fig. 3B). Untagged protein was not detected in reactions that included tmRNA-SmpB, even after addition of puromycin to release any peptidyl-tRNA that remained in the ribosome, indicating that *trans*-translation was efficient and little peptidyl-tRNA remained in the ribosome. If the ribosome translates to within 8 codons of the PAM site as observed *in vivo*, a protein of 16.9 kDa should be produced. To estimate the location of the tagging site, we made a 153 amino acid protein derived from DHFR. Reactions using this template produced proteins with similar gel mobility to the truncated proteins generated by the dCas9 block, suggesting that translation extends to within ∼8 codons of the PAM site in a blocked transcription-translation complex *in vitro* (Fig. 3B) as well as *in vivo* (Fig. 2B). These results demonstrate that in a purified *in vitro* system, ribosomes behind a blocked RNA polymerase can translate to the 3’ end of the mRNA, where they are rescued by *trans*-translation.

**Fig 3.**
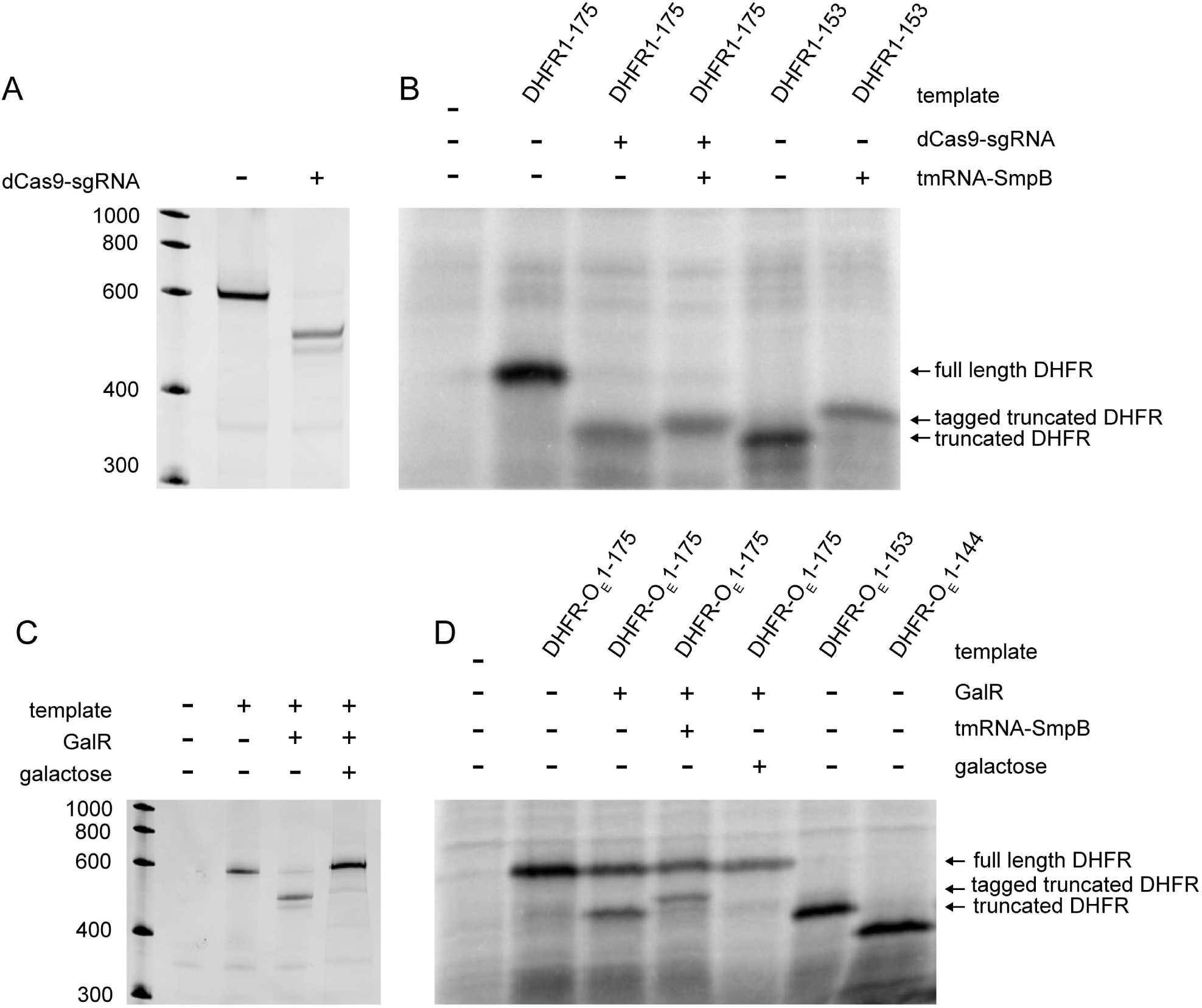
Outcomes from blocking RNA polymerase with a DNA-binding protein *in vitro.* (*A*) *In vitro* transcription with or without dCas9-sgRNA. DNA encoding DHFR was incubated with or without dCas9-sgRNA and used as template for *in vitro* transcription with *E. coli* RNA polymerase. Transcripts were separated by denaturing urea-PAGE and detected by ethidium bromide staining. (*B*) *In vitro* transcription-translation and *trans*-translation reactions with or without dCas9-sgRNA. DNA template encoding full-length DHFR (residues 1-175) was incubated with or without dCas9-sgRNA and transcribed and translated *in vitro* in the absence or presence of tmRNA-SmpB. Newly synthesized proteins were detected by incorporation of ^35^S-methionine and phosphorimaging. Control reactions used truncated DHFR template (residues 1-153) that did not contain a stop codon. Products corresponding to full length DHFR, tagged truncated DHFR, and truncated DHFR are indicated. (*C*) *In vitro* transcription of DNA template encoding DHFR with a GalR operator site (residues 1-175). (*D*) *In vitro* translation and *trans*-translation reactions with or without GalR. Control reactions used truncated DHFR template (residues 1-153 or residues 1-144). Representative images from at least three biological repeats are shown.

To ensure that the ribosome rescue mechanism we observed was not unique to complexes in which RNA polymerase was blocked by dCas9, we repeated the *in vitro* experiment using the *E. coli* galactose operon repressor protein, GalR. GalR binds to the 20-mer exterior operator (O_E_) site with K_d_ = 4 nM and this binding is sufficient to block transcription from the *P*1 promoter of the *gal* operon *in vitro* (34). The affinity of GalR for DNA is reduced by binding with D-galactose (34,35). We cloned the O_E_ site within the coding sequence of a DHFR template DNA and used it for transcription, translation, and *trans*-translation assays. Inclusion of GalR in reactions with this template resulted in truncated transcript and truncated protein (Fig. 3C & D). When tmRNA-SmpB were also included in translation reactions, the truncated protein was tagged and little untagged truncated protein was observed. Truncation and tagging were not observed when galactose was added to the reaction, consistent with galactose preventing GalR from binding DNA. A DHFR template corresponding to residues 1-153 produced a protein with gel mobility similar to the truncated protein produced by the GalR block, suggesting that translation extends to within 3 codons of the O_E_ site (Fig. 3D). These data indicate that the ribosome translates mRNA that would be situated within RNA polymerase. By contrast, the truncated protein made by a GalR block was of a slower mobility than a template truncated to residues 1-144, which would be the expected site of the blocked ribosome if it stopped at the edge of RNA polymerase. The major difference between reactions with GalR and dCas9 was the increased amount of full-length protein in the GalR reactions. It is possible that GalR has a lower affinity for DNA in the *in vitro* translation buffer than dCas9, resulting in more unbound DNA and more full-length transcript. Alternatively, weaker binding by GalR may allow RNA polymerase to push through the block more frequently. Despite this difference, these experiments show that an endogenous DNA-binding protein like GalR can also block transcription-translation complexes, and ribosomes from these complexes are rescued by *trans*-translation.

Translation to the 3’ end of the transcript in a blocked transcription-translation complex indicates that the nascent transcript is released from RNA polymerase with the ribosome still engaged. The translating ribosome could either trigger release of the mRNA from RNA polymerase without removing RNA polymerase from the DNA, or RNA polymerase could dissociate from the DNA and release the transcript. To distinguish between these possibilities, we tested whether RNA polymerase and the nascent transcript remain associated with DNA after collision of a translating ribosome. We first set up *in vitro* transcription-translation reactions with 5’ biotinylated DNA template in the presence or absence of dCas9-sgRNA and used streptavidin-coated magnetic beads to isolate the DNA and any bound proteins. 73 ± 22% of RNA polymerase remained associated with the DNA at the end of the transcription-translation reaction (Fig. 4A). When transcription was blocked by GalR, 41 ± 16% of RNA polymerase remained associated with the biotinylated DNA (Fig. 4B). These results indicate that the ribosome translates to the 3’ end of the mRNA while RNA polymerase remains bound to the DNA. The higher percentage of RNA polymerase that is released in the reactions with GalR compared to the reactions with dCas9 is likely due to higher amount of unblocked RNA polymerase.

**Fig. 4.**
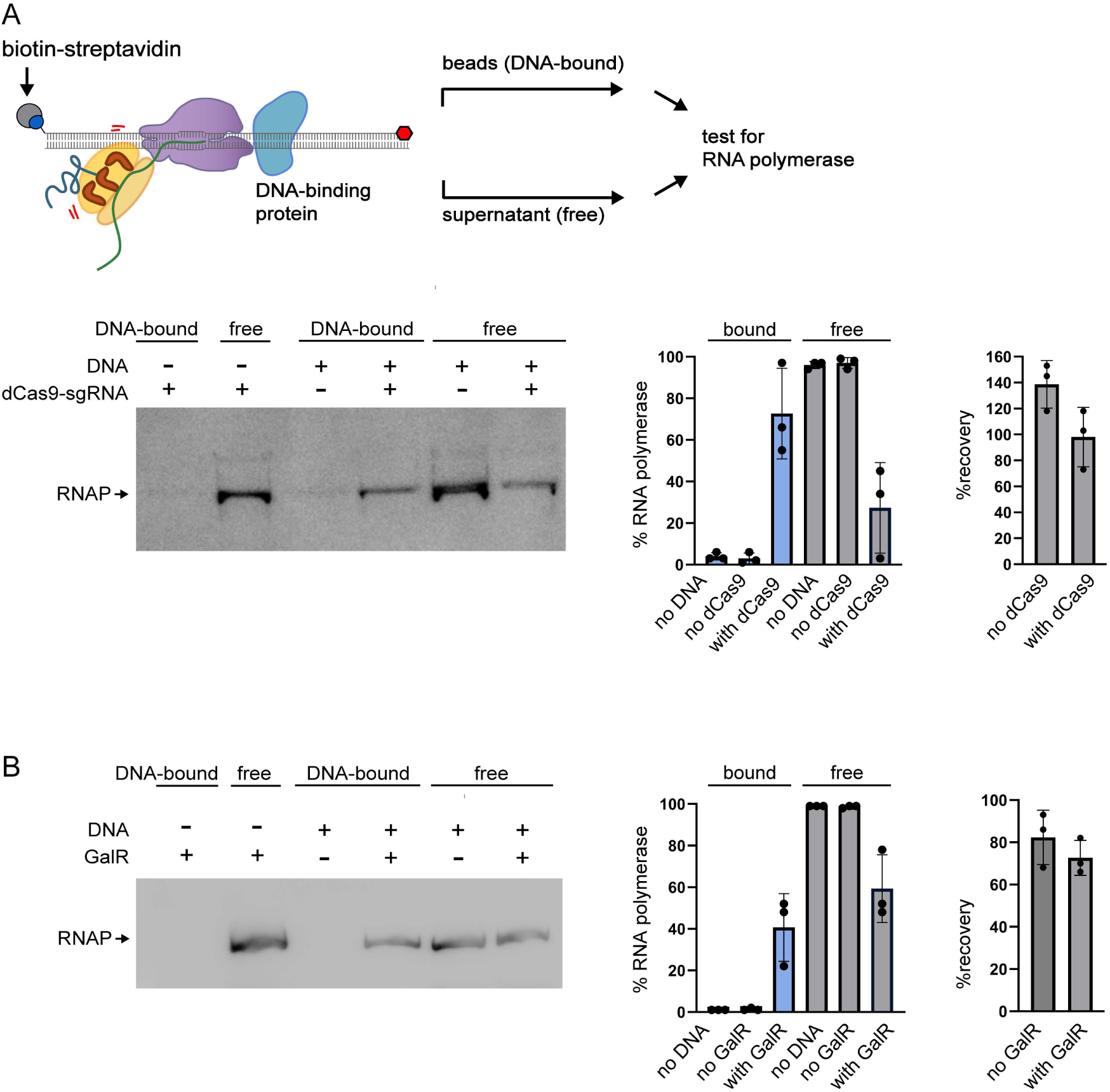
RNA polymerase remains associated with DNA at a protein block. Biotinylated DNA encoding DHFR was incubated with or without dCas9-sgRNA (*A*) or GalR (*B*) and used as template for *in vitro* transcription-translation. The DNA was pulled down with streptavidin beads and the amount of RNA polymerase in the DNA-bound and free fractions were measured by western blotting. The percentage of RNA polymerase in each fraction was calculated and the mean ± standard deviation from three biological repeats is shown. The recovery of RNA polymerase for the combined DNA-bound and free fractions in each sample is shown relative to a control reaction containing no DNA template.

To confirm that the ribosome triggers release of mRNA from RNA polymerase, we set up transcription reactions using biotinylated template DNA and used streptavidin beads to pull down the DNA and any factors that remained bound to the DNA after the reaction (Fig. 5A). When sgRNA-dCas9 was included in the reaction, 87 ± 7% of the transcript was pulled down with the biotinylated DNA, indicating that it was still associated with RNA polymerase. To measure transcript release, we set up transcription-translation reactions in the presence of α[^32^P]-UTP and purified the blocked translation complex with streptavidin beads before initiating translation. Because the translation reaction includes many purified factors that could, in principle, affect release of the mRNA, we used a mock translation reaction in which ATP and GTP were omitted as a negative control. When translation was active, 66 ± 18% of the transcript was released (Fig. 5B). These data demonstrate that translation removes the nascent transcript from blocked transcription-translation complexes, consistent with possibility 4 from Figure 1.

**Fig. 5.**
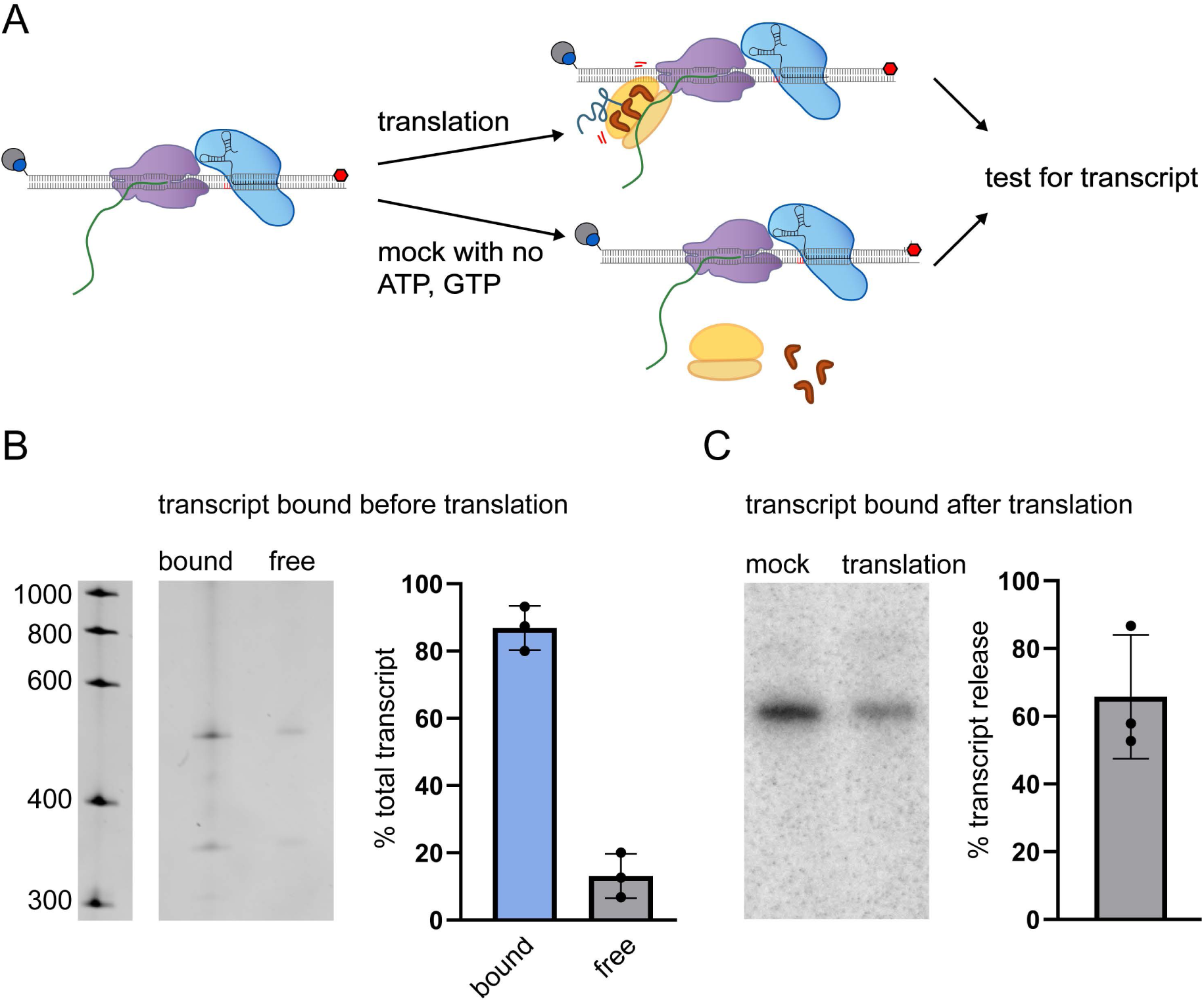
Translating ribosomes release the transcript from blocked RNA polymerase. (*A*) Biotinylated DNA encoding DHFR was incubated with dCas9-sgRNA and used as template for *in vitro* transcription. Transcripts from *in vitro* transcription were pulled out with streptavidin beads, separated by denaturing urea-PAGE, and detected by ethidium bromide staining. The mean ± standard deviation from 3 biological repeats is shown. (*B*) Biotinylated DNA encoding DHFR was incubated with dCas9-sgRNA and used as template for *in vitro* transcription in the presence of [α-^32^P]UTP. The DNA and bound RNA polymerase and transcript were pulled out with streptavidin beads and used as template for *in vitro* translation. A control reaction contained all translation components except ATP and GTP. The amount of transcript that remained associated with the DNA was separated on urea-polyacrylamide gels and quantified by phosphorimaging. The percentage of the transcript that was released from RNA polymerase was calculated and the mean ± standard deviation from 3 biological repeats is shown.

## Discussion

Our results suggest a model for rescuing all the components of a transcription-translation complex that is blocked by a DNA-binding protein or DNA damage. The leading ribosome initiates rescue by triggering release of the nascent transcript from RNA polymerase. All ribosomes translating that transcript will then translate to the 3’ terminus of the mRNA and can be rescued by *trans*-translation, leading to recycling of the ribosome, proteolysis of the nascent polypeptide, and degradation of the mRNA (Possibility 4 in Figure 1). *In vitro,* RNA polymerase remains on the DNA after the transcript is removed. In the cell, this post-termination RNA polymerase is likely to be available for recycling by RapA or other factors (36). Our results are consistent with previous work showing that a translating ribosome can destroy an RNA polymerase elongation complex stalled at DNA lesions (3). Our data show that a translating ribosome can also act on RNA polymerase stalled by a DNA-binding protein, demonstrate the fate of the translating ribosomes, and provide a mechanism for destroying the elongation complex by extracting the mRNA.

Optical trapping experiments to measure the force generated by a translating ribosome (∼13 ± 2 pN) (37) and the force required to pull the transcript out of RNA polymerase (>30 pN) (38) suggest that a translating ribosome is unlikely to exert enough force to mechanically extract mRNA from a transcribing RNA polymerase. It is possible that the transcript is bound less tightly by a blocked RNA polymerase, so the ribosome can simply pull it out. Alternatively, the collision between the translating ribosome and the blocked RNA polymerase may decrease the stability of the RNA polymerase elongation complex, for example through a structural change in RNA polymerase.

Bacterial transcription factors frequently bind within protein coding sequences (39,40), and are therefore likely to lead to blocked transcription-translation complexes that need to be rescued. For example, LacI binds the *lac* operator with sub-nanomolar affinity (41), stronger than GalR or dCas9. The *lac* operator overlaps with the *lacI* coding sequence, and binding of LacI to the *lac* operator leads to a dramatic increase in *trans*-translation on newly synthesized LacI (42). Although RNA polymerase stalling at these locations has not been tested directly, the observed *trans*-translation on LacI is consistent with the model presented here. Genome-wide CHIP-seq identification of transcription factor binding sites in *E. coli* reveals transcription factor binding across a substantial fraction of the genome (40,43). In principle, proteins bound within a coding region and ≥75 bases from the start codon could lead to blocked transcription-translation complexes because blocking RNA polymerase at these locations would result in a nascent transcript long enough for a ribosome to initiate translation. Such binding sites are abundant. In fact, one CHIP-seq dataset where transcription factors were expressed from their native promoters showed transcription factor binding in a location that could lead to a blocked transcription-translation complex in 610 genes (>13% of all genes in the genome) (40). The chance that blocked transcription-translation complexes will be generated at one of these sites will depend on the concentration of the transcription factor in the cell, but there are enough sites to account for a large fraction of the observed *trans*-translation activity.

Our model for the rescue of blocked transcription-translation complexes can also explain published data demonstrating that mutants incapable of *trans*-translation are hypersensitive to aza-C-induced DNA-protein crosslinks and streptolydigin, an antibiotic that blocks RNA polymerase elongation (44–46). Translation of the nascent transcript on an RNA polymerase blocked by either DNA-protein crosslinks or streptolydigin would release the mRNA, and the ribosomes would translate to the 3’ end. In wild-type cells, these ribosomes would be rescued by *trans*-translation, but mutants lacking *trans*-translation would have to use alternative ribosome rescue pathways that have lower capacity and generate proteotoxic stress, making them more sensitive to the blocking agents.

Notably, our proposed mechanism for rescuing blocked transcription-translation complexes does not require any new activity for tmRNA-SmpB. Because the ribosome translates to the 3’ end of the mRNA before *trans*-translation, tmRNA-SmpB can access the empty A site and mRNA channel as observed for all known *trans*-translation substrates *in vitro* and *in vivo* (5).

## Materials and methods

### Bacterial Strains and Plasmids

Strains are described in Table 1 and were grown in lysogeny broth at 37°C with ampicillin (50 µg/mL), chloramphenicol (20 µg/mL), kanamycin (30 µg/mL), or spectinomycin (50 µg/mL), where appropriate. DNA oligos are described in Table 2. All DNA oligos were synthesized by Integrated DNA Technologies (Coralville, IA).

**Table 1.** Bacterial strains and plasmids.

| Name | Description | Source or reference |
| --- | --- | --- |
| **Strain or plasmid** |  |  |
| KCK 629 | DH10B with pCA24N-DHFRmet7; Cm<sup>R</sup> | This study |
| KCK 638 | DH5α pCA24N-DHFRmet7-galR_OE; Cm<sup>R</sup> | This study |
| KCK 630 | MRE600 | (47) |
| KCK 186 | MG1655Δ*ssrA*::cat | This study |
| KCK 631 | BL21(DE3) with pET28b-dCas9; Kan<sup>R</sup> | This study |
| KCK 639 | BL21(DE3) pET28b-galR; Kan<sup>R</sup> | This study |
| NSR 4 | BL21(DE3) with pET28b-SmpB | (48) |
| KCK 349 | X90*ssrAH6D* | (25) |
| KCK 632 | X90Δ*ssrA*::cat | This study |
| KCK 633 | X90*ssrAH6D* pBAD-dCas9,sgRNA; pTrc-FLAGEGFP | This study |
| KCK 634 | X90Δ*ssrA*::cat pBAD-dCas9,sgRNA; pTrc-FLAGEGFP | This study |
| KCK 635 | DH5α pBAD-dCas9,sgRNA; pTrc99a-EGFP_stop | This study |
| KCK 636 | DH5α pBAD-dCas9,sgRNA; pTrc99a-EGFP_H6D | This study |
| KCK 637 | X90*ssrAH6D* pBAD-dCas9,sgRNA; pTrc99a-EGFP_TTT | This study |
| JW0047 | pCA24N-*folA*; DHFR | (49) |
| pET28b | For expression from the T7 promoter | Novagen |
| pEcgRNA | Contains the gRNA scaffold | Addgene<br>166581 |
| pJMP1337 | Contains dCas9 | (50) |
| pBAD24 | Contains an arabinose inducible promoter | (51) |
| pACYC184 | Contains the p15A origin of replication | (52) |
| pEGFP-N2 | Contains EGFP | Takara<br>Bio |
| pTrc99a | Contains the *trc* promoter | (53) |
| JW2667 | AG1 with pCA24N-*alaS* | (49) |
| JW1865 | AG1 with pCA24N-*argS* | (49) |
| JW0913 | AG1 with pCA24N-*asnS* | (49) |
| JW1855 | AG1 with pCA24N-*aspS* | (49) |
| JW0515 | AG1 with pCA24N-*cysS* | (49) |
| JW0666 | AG1 with pCA24N-*glnS* | (49) |
| JW2395 | AG1 with pCA24N-*gltx* | (49) |
| NM 3-39 | DH5α with pCA24N-*glyQ-glyS* | This<br>study |
| JW2498 | AG1 with pCA24N-*hisS* | (49) |
| JW0024 | AG1 with pCA24N-*ileS* | (49) |
| JW0637 | AG1 with pCA24N-*leuS* | (49) |
| JW4090 | AG1 with pCA24N-*lysU* | (49) |
| JW2101 | AG1 with pCA24N-*metG* | (49) |
| NM 3-45 | DH5α with pCA24N-*pheS-pheT* | This<br>study |
| JW0190 | AG1 with pCA24N-*proS* | (49) |
| JW0876 | AG1 with pCA24N-*serS* | (49) |
| JW1709 | AG1 with pCA24N-*thrS* | (49) |
| JW3347 | AG1 with pCA24N-*trpS* | (49) |
| JW1629 | AG1 with pCA24N-*tyrS* | (49) |
| JW4215 | AG1 with pCA24N-*valS* | (49) |
| JW3249 | AG1 with pCA24N-*fmt* | (49) |
| JW2502 | AG1 with pCA24N-*ndk* | (49) |
| JW0867 | AG1 with pCA24N-*infA* | (49) |
| JW3137 | AG1 with pCA24N-*infB* | (49) |
| JW5829 | AG1 with pCA24N-*infC* | (49) |
| JW1202 | AG1 with pCA24N-*prfA* | (49) |
| JW5847 | AG1 with pCA24N-*prfB* | (49) |
| JW5873 | AG1 with pCA24N-*prfC* | (49) |
| JW0167 | AG1 with pCA24N-*frr* | (49) |
| JW3302 | AG1 with pCA24N-*fusA* | (49) |
| JW0165 | AG1 with pCA24N-*tsf* | (49) |
| JW3301 | AG1 with pCA24N-*tufA* | (49) |

**Table 2.**
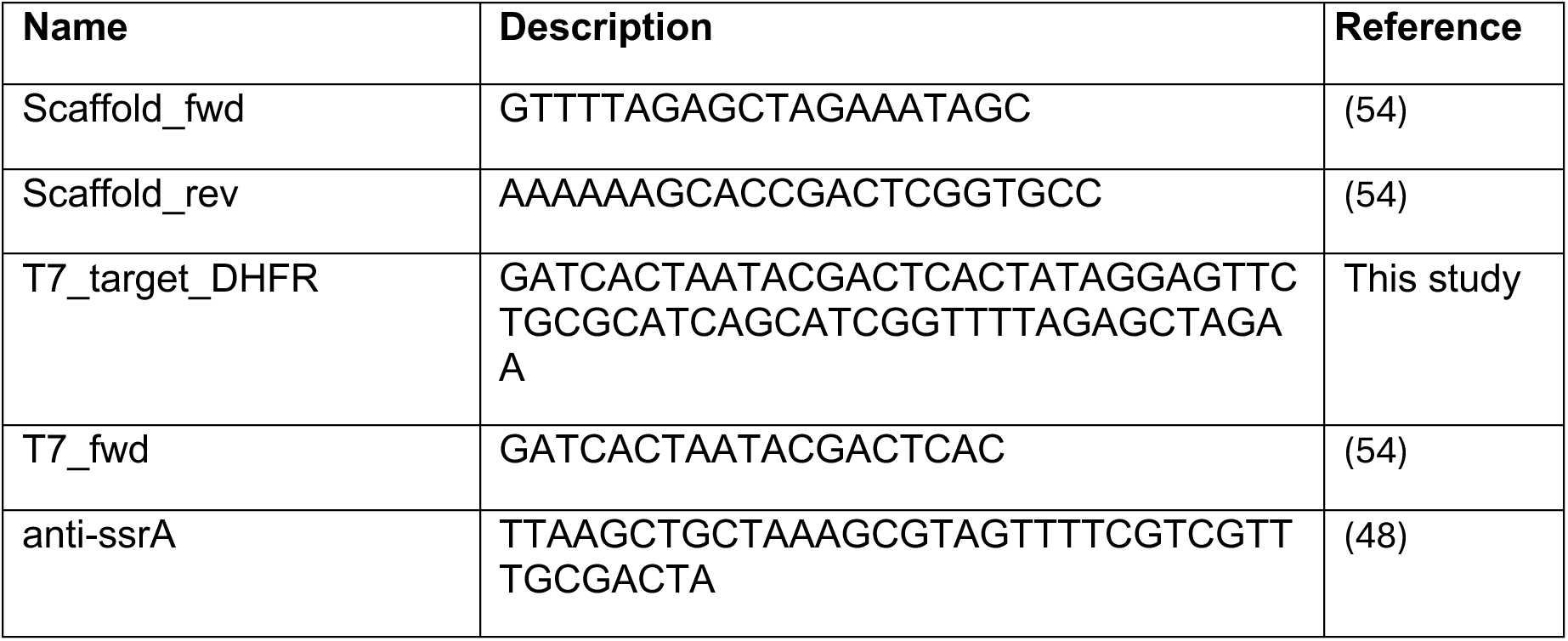
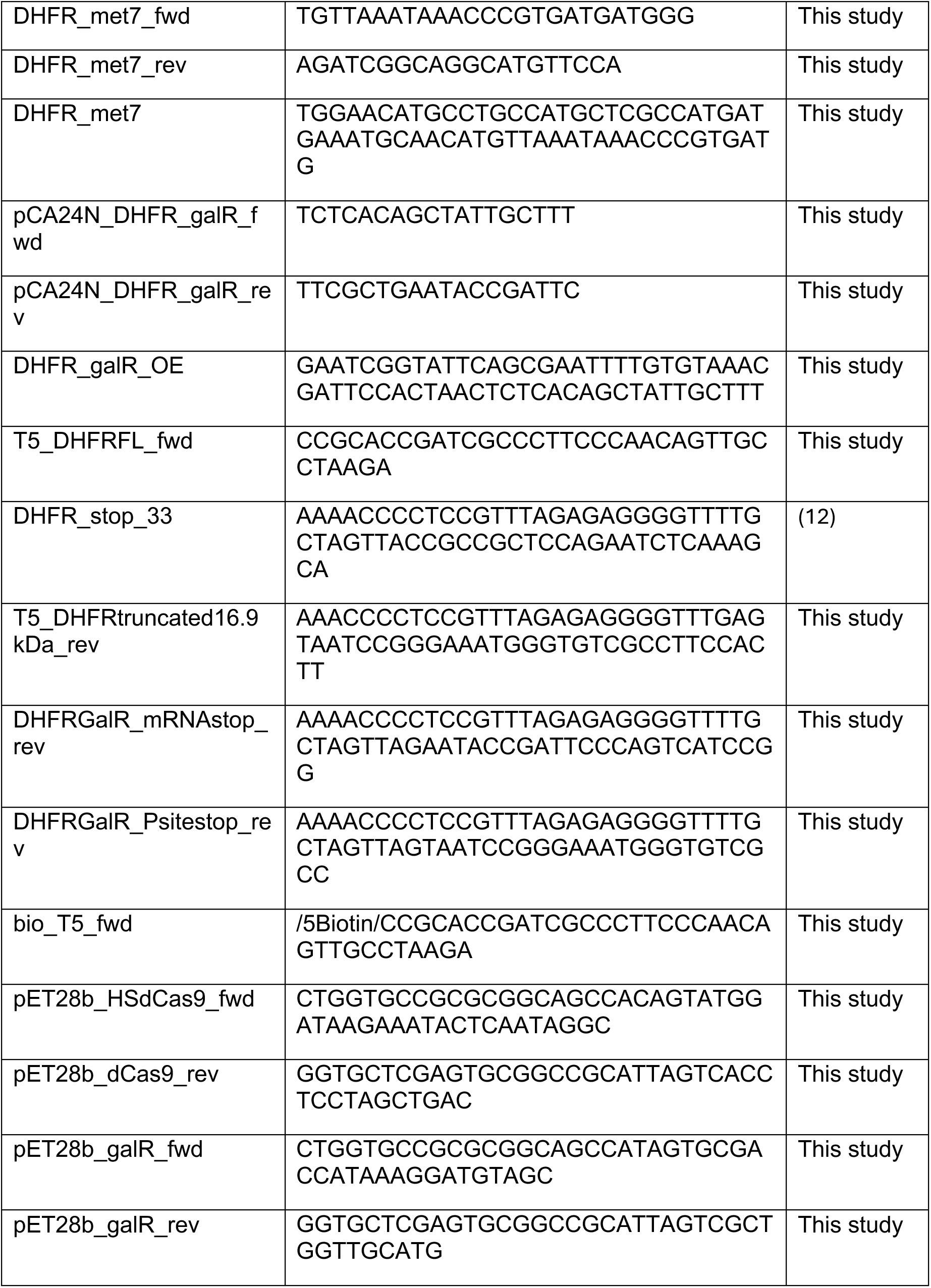
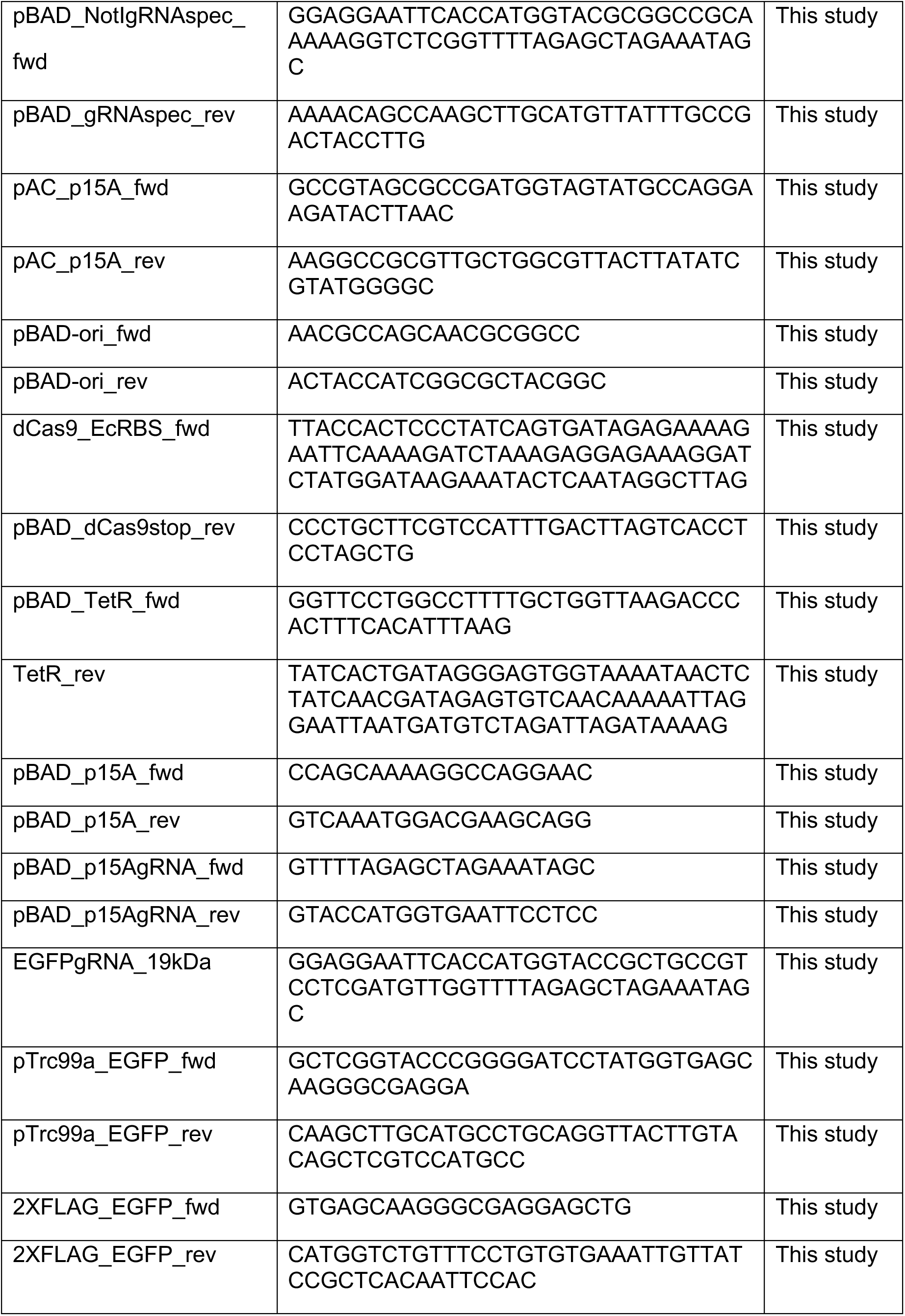
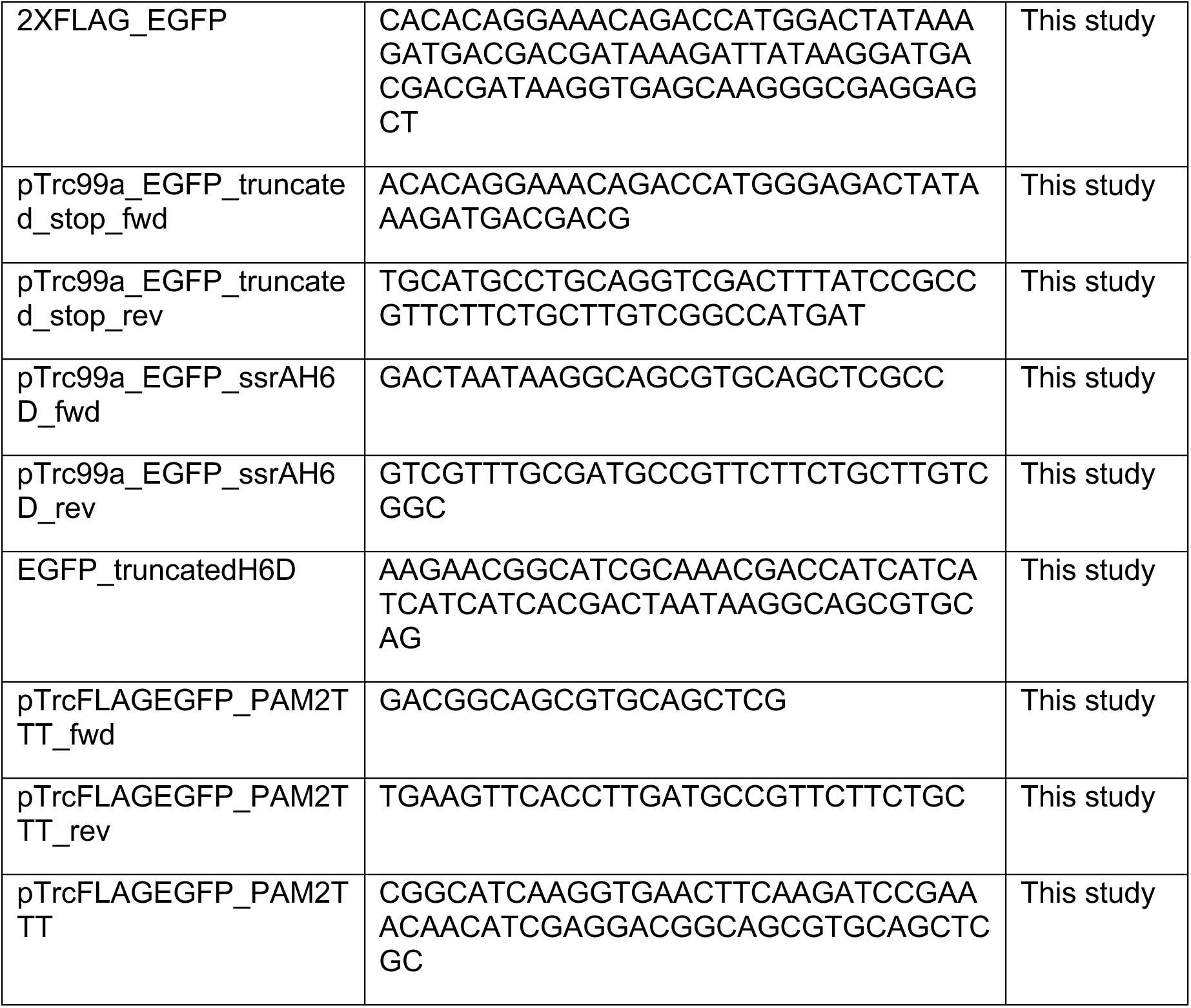
Oligonucleotides.

To construct pCA24N-DHFRmet7 containing 7 additional methionines to DHFR, pCA24N-folA was PCR-amplified with primers DHFR_met7_fwd and DHFR_met7_rev, digested with *Dra*I, and assembled by HiFi DNA assembly (New England Biolabs) with the ssDNA oligo DHFR_met7.

To construct pCA24N-DHFR-GalR_OE with the GalR operator sequence replacing bases 420-439 of the *folA* (DHFR) gene, pCA24N-DHFRmet7 was PCR-amplified with primers pCA24N_DHFR_galR_fwd and pCA24N_DHFR_galR_rev, digested with *Dpn*I, and assembled by HiFi DNA assembly with the ssDNA oligo DHFR_galR_OE containing the 20-mer GalR operator sequence, TTGTGTAAACGATTCCACTA.

To construct pET28b-dCas9, the dCas9 gene was PCR-amplified from pJMP1337 (50) with primers pET28b_HSdCas9_fwd and pET28b_dCas9_rev and assembled by HiFi DNA assembly into pET28b (Novagen; Merck KGaA, Darmstadt, Germany) cut with *Hind*III and *Nde*I. The resulting plasmid was transformed into BL21(DE3) (Novagen).

To construct pET28b-galR, the *galR* gene was PCR-amplified from MG1655 genomic DNA with primers pET28b_galR_fwd and pET28b_galR_rev and assembled into pET28b cut with *Hind*III and *Nde*I. The resulting plasmid was transformed into BL21(DE3) (Novagen).

To construct pBAD-p15A-tet-dCas9-GFP-gRNA-spec, pEcgRNA was PCR-amplified with primers pBAD_NotIgRNAspec_fwd and pBAD_gRNAspec_rev and assembled into pBAD24 (51) cut with *Kpn*I and *Pae*I to generate pBAD-*Not*I-gRNA-spec. The p15A origin was amplified from pACYC184 (52) with primers pAC_p15A_fwd and pAC_p15A_rev and assembled with a *Sca*I-digested PCR product amplified off of pBAD-*Not*I-gRNA-spec with primers pBAD-ori_fwd and pBAD-ori_rev to generate pBAD-p15A-gRNA-spec. An N20 sequence targeting EGFP was designed with the CRISPy-web 2.0 tool (55) and was added by assembling EGFPgRNA_19kDa ssDNA oligo with a *Not*I-digested PCR product amplified off of pBAD-p15A-gRNA-spec with primers pBAD_p15AgRNA_fwd and pBAD_p15AgRNA_rev to generate pBAD-p15A-GFP-gRNA.

dCas9 and TetR were amplified off of pJMP1337 with dCas9_EcRBS_fwd and pBAD_dCas9stop_rev and pBAD_TetR_fwd and TetR_rev, respectively, and assembled with *Bsu*15I-digested PCR product amplified off of pBAD-p15A-GFP-gRNA with pBAD_p15A_fwd and pBAD_p15A_rev to generate pBAD-p15A-tet-dCas9-GFP-gRNA-spec.

To construct pTrc99a-2XFLAG-EGFP, EGFP was amplified from pEGFP-N2 (Takara Bio USA, San Jose, CA) with primers pTrc99a_EGFP_fwd and pTrc99a_EGFP_rev and assembled into pTrc99a (53) cut with *Sal*I and *Xba*I to generate pTrc99a-EGFP. Tandem FLAG tags were added with 2XFLAG_EGFP ssDNA oligo assembled with *Dpn*I-digested PCR product amplified off of pTrc99a-EGFP with primers 2XFLAG_EGFP_fwd and 2XFLAG_EGFP_rev. Truncated EGFP with a stop codon was constructed by PCR off pTrc99a-2XFLAG-EGFP with primers pTrc99a_EGFP_truncated_stop_fwd and pTrc99a_EGFP_truncated_stop_rev and assembled into pTrc99a cut with *Sal*I and *Xba*I to generate pTrc99a-EGFP_stop. Truncated EGFP with HHHHHHD added to the C-terminal end was generated by assembling EGFP_truncatedH6D ssDNA oligo with a *Dpn*I-digested PCR product amplified off of pTrc99a-2XFLAGEGFP with primers pTrc99a_EGFP_ssrAH6D_fwd and pTrc99a_EGFP_ssrAH6D_rev to obtain pTrc99a-EGFP_H6D. EGFP with the PAM site mutated to TTT was constructed by assembling the ssDNA oligo pTrcFLAGEGFP_PAM2TTT into a *Dpn*I-digested PCR product amplified from pTrc99a-2XFLAG-EGFP with primers pTrcFLAGEGFP_PAM2TTT_fwd and pTrcFLAGEGFP_PAM2TTT_rev to generate pTrc99a-EGFP_TTT.

MG1655Δ*ssrA*::*cat* was constructed by λ Red recombination (56) into MG1655.

X90Δ*ssrA*::*cat* was constructed by P1 transduction (57) of MG1655Δ*ssrA*::*cat* into the X90 background.

### Purification of components for *in vitro* transcription and translation

*E. coli* translation factors were purified based on the OnePot PURE system (58) with the following modifications. All translation factors were expressed from the ASKA collection (49) with 1 mM isopropyl-thio-β-D-galactoside (IPTG) for 4 h. *E. coli* release factors RF1, RF2, and RF3 from the ASKA collection were expressed and purified separately using the same buffers as the OnePot PURE system. *E. coli* EF-Tu was purified as described (59). Because the complete gene sequences of the aminoacyl-tRNA synthetases for glycine and phenylalanine are split between two gene annotations in the ASKA collection, we cloned GlyRS and PheRS individually into pCA24N. Components not available from the ASKA collection were purchased: creatine kinase (Roche, Indianapolis, IN; cat. 10127566001), myokinase (Sigma-Aldrich, Burlington, MA; cat. M3003), thermostabile inorganic pyrophosphatase (New England Biolabs, Ispwich, MA; cat. M0296), and *E. coli* RNAP holoenzyme (NEB, cat. M0551).

*E. coli* tRNAs were purified from MRE600 as described (60). The energy solution containing tRNAs was prepared as described (58) except where indicated.

Ribosomes were purified from MG1655Δ*ssrA*::*cat* by centrifugation through a sucrose cushion. Cells were grown to OD_600_ ∼0.7, harvested by centrifugation at 6,953 *g* for 10 min, and resuspended in ribosome resuspension buffer (20 mM HEPES [pH 7.5 with KOH], 60 mM NH_4_Cl, 12 mM MgCl_2_, 0.5 mM EDTA, 6 mM β-mercaptoethanol (β-ME)). Cells were lysed by sonication and the lysate was cleared by centrifugation at 28,000 *g* for 20 min at 4°C. Crude ribosomes were pelleted over ribosome sucrose cushion buffer (20 mM HEPES-KOH [pH 7.5], 500 mM NH_4_Cl, 10 mM MgCl_2_, 0.5 mM EDTA, 37% (w/v) sucrose, 6 mM β-ME) at 257,000 *g* for 2 h at 4°C. Ribosomes were washed and resuspended in ribosome resuspension buffer to a final concentration of ∼13.5 µM.

Full-length DNA template for *in vitro* transcription and translation was prepared from pCA24N-DHFRmet7 or pCA24N-DHFR-GalR_OE with primers T5_DHFRFL_fwd and DHFR_stop_33. Truncated template without a stop codon was prepared from pCA24N-DHFRmet7 with primers T5_DHFRFL_fwd and T5_DHFRtruncated16.9kDa_rev. Truncated template corresponding to 17.2 kDa was prepared from pCA24N-DHFR-GalR_OE with primers T5_DHFRFL_fwd and DHFRGalR_mRNAstop_rev. Truncated template corresponding to 16.1 kDa was prepared from pCA24N-DHFR-GalR_OE with primers T5_DHFRFL_fwd and DHFRGalR_Psitestop_rev. Biotinylated DNA template for *in vitro* transcription and translation was made from pCA24N-DHFRmet7 or pCA24N-DHFR-GalR_OE with primers bio_T5_fwd and DHFR_stop_33.

dCas9 was purified as described (5) except the protein was eluted in 500 mM imidazole and stored in 220 mM KCl, 20 mM HEPES, [pH 7.5] (21). sgRNA was purified as described (54) except sgRNA scaffold was PCR amplified from pEcgRNA and overlapping PCR was performed with the ssDNA oligo T7_target_DHFR (Table 2).

GalR was purified based on (61) except where indicated. BL21(DE3) pET28b-GalR was grown in terrific broth (24 g/L yeast extract, 20 g/L tryptone, 4 mL/L glycerol, 0.017 M KH_2_PO_4_, 0.072 M K_2_HPO_4_) at 37°C to OD_600_ ∼ 0.7 and induced with 0.2 mM IPTG at 18°C for 16 h. Cells were harvested by centrifugation at 6,953 *g* for 10 min and resuspended in lysis buffer (25 mM Tris [pH 8.0], 5 mM EDTA, 600 mM KCl, 1 mM DTT, 10 mM imidazole, 1 mg/mL lysozyme) on ice for 2 h. Cells were lysed in an LM20 microfluidizer processor (Microfluidics, Westwood, MA) at 20,000 psi and cell debris were removed by centrifugation at 28,000 *g* for 20 min at 4°C. Cleared lysate was bound in batch to HisPur Ni-NTA agarose resin (Thermo Fisher Scientific, Waltham, MA) for 1 h at 4 °C, washed 4 times with wash buffer (25 mM Tris [pH 8.0], 5 mM EDTA, 600 mM KCl, 1 mM DTT, 10 mM imidazole) and eluted in elution buffer (25 mM Tris [pH 8.0], 1 mM EDTA, 600 mM KCl, 1 mM DTT, 500 mM imidazole, 15% (v/v) glycerol). Fractions containing GalR were dialyzed in storage buffer (25 mM Tris [pH 8.0], 1 mM EDTA, 600 mM KCl, 1 mM DTT, 15% (v/v) glycerol) at 4 °C. Dialyzed protein was concentrated to ∼82 µM and frozen at -20°C in storage buffer plus 50% (v/v) glycerol.

tmRNA was prepared by *in vitro* transcription and purified by UV shadowing as described (48). For purification of SmpB BL21(DE3) pET28b-SmpB was grown in terrific broth at 37°C to OD_600_ ∼ 0.6 and induced with 1 mM IPTG for 3 h. Cells were harvested by centrifugation at 6,953 *g* for 10 min and resuspended in buffer A (100 mM NaH_2_PO_4_ [pH 7.6], 150 mM KCl, 6 M GuHCl, 10 mM imidazole). Cells were lysed by sonication and cell debris were removed by centrifugation at 28,000 *g* for 20 min at 4°C. Cleared lysate was bound in batch to HisPur Ni-NTA agarose resin (Thermo Fisher Scientific) for 1 h at 4 °C, washed 4 times with buffer A, and eluted in elution buffer (100 mM NaH_2_PO_4_ [pH 5.9], 300 mM KCl, 6 M GuHCl, 1 M imidazole). Fractions containing SmpB were dialyzed against storage buffer (50 mM HEPES-KOH [pH 7.6], 300 mM KCl, 10 mM MgCl_2_, 20% (v/v) glycerol, and 7 mM β-ME) at 4 °C. Dialyzed protein was concentrated to ∼20 µM and frozen at -80°C in storage buffer.

### *In vitro* transcription

dCas9 blocks were assembled by incubating 245 nM DNA template encoding DHFR with 1.22 µM dCas9 and 1.22 µM sgRNA in dCas9 reaction buffer (20 mM Tris [pH 7.5], 100 mM KCl, 5 mM MgCl_2_, 1 mM DTT, 5% (v/v) glycerol) for 20 min at 37°C. DNA template with or without dCas9 was added to energy solution (32.5 mM HEPES [pH 7.5], 7.7 mM magnesium acetate, 65 mM potassium glutamate, 0.65 mM dithiothreitol (DTT), 13 mM creatine phosphate, 13 µM folinic acid, 1.3 mM spermidine, 1.3 mM ATP, 1.3 mM GTP, 0.65 mM CTP, 0.65 mM UTP, amino acid mix (0.13 mM methionine and the other 19 common amino acids at 1.3 mM), 1.7 mg/mL tRNA) containing 312 nM *E. coli* RNA polymerase holoenzyme to a final DNA template concentration of 143 nM. Reactions were incubated for 5 min at 37°C, DNase I treated, heated to 65°C in urea loading buffer (8 M urea, 5 mM Tris [pH 7.5], 25 mM EDTA, 0.05% bromophenol blue, 0.05% xylene cyanol), and resolved on 5% denaturing urea-PAGE. Transcripts were visualized by ethidium bromide staining and imaged on a ChemiDoc MP (Bio-Rad, Hercules, CA).

GalR blocks were assembled by incubating 165 nM DNA template encoding DHFR with the GalR operator site with 16 µM GalR in energy solution for 20 min at 37°C. For reactions with galactose, 16 µM GalR was incubated with 1.1 M galactose in energy solution for 10 min at 37°C, followed by addition of 165 nM DNA template encoding DHFR with the GalR operator site. Transcription was initiated with 292 nM *E. coli* RNA polymerase holoenzyme to a final DNA template concentration of 143 nM. Reactions were incubated for 5 min at 37°C, DNase I treated, heated to 65°C in urea loading buffer, and resolved on 5% denaturing urea-PAGE. Transcripts were visualized by ethidium bromide staining and imaged on a ChemiDoc MP (Bio-Rad).

### *In vitro* transcription and translation and *trans*-translation

dCas9 blocks were assembled by incubating 140 nM DNA template encoding DHFR with 1.2 µM dCas9 and 1.2 µM sgRNA at 37°C for 20 min. *In vitro* transcription and translation were initiated by addition of energy solution (at the concentrations indicated for *in vitro* transcription), OnePot translation factor mix (1.7 mg/mL), *E. coli* RNA polymerase holoenzyme (100 nM), release factor mix containing RF1, RF2, and RF3 (23 ng/µL), EF-Tu (10 µM), ribosomes (1.4 μM), and [^35^S]-methionine (1 µCi, 1175 Ci/mmol) (Revvity, Waltham, MA), yielding a final DNA template concentration of 34 nM.

GalR blocks were assembled by incubating 96 nM DNA template encoding DHFR containing the GalR operator site with 8.5 µM GalR in energy solution 37°C for 20 min. For reactions with galactose, 8.5 µM GalR was incubated with 0.6 M galactose in energy solution for 10 min at 37°C, followed by addition of 96 nM DNA template encoding DHFR with the GalR operator site. *In vitro* transcription and translation were initiated by addition of OnePot translation factor mix (1.7 mg/mL), *E. coli* RNA polymerase holoenzyme (100 nM), release factor mix containing RF1, RF2, and RF3 (23 ng/µL), EF-Tu (10 µM), ribosomes (1.4 μM), and [^35^S]-methionine (1 µCi, 1175 Ci/mmol) (Revvity), yielding a final DNA template concentration of 44 nM. *trans*-Translation reactions for dCas9 and GalR were set up in the same manner but included 1.4 µM tmRNA and 1.4 µM SmpB. Reactions without tmRNA-SmpB were supplemented with 0.5 µM anti-SsrA oligonucleotide to limit *trans*-translation activity (48). Reactions were incubated at 37°C for 20 min, and 3.3 µg/µL puromycin was added to hydrolyze any remaining peptidyl-tRNA. Proteins were acetone precipitated, heated to 95°C in Laemmli sample buffer (100 mM Tris [pH 6.8], 2% SDS, 20% glycerol, 0.001% bromophenol blue, 6 mM β-ME), resolved on 15% SDS-PAGE, and visualized by phosphorimaging in a Typhoon RGB (Cytiva, Wilmington, DE). Band intensities were quantified in ImageJ version 1.54g (62).

### Measurement of RNA polymerase association with DNA

We validated the efficiency of pulling out biotinylated DNA in our *in vitro* experimental set-up by incubating 30 nM of 5’ biotinylated DHFR DNA template or non-biotinylated DNA with Dynabeads MyOne Streptavidin C1 magnetic beads (Invitrogen, Waltham, MA) in a buffer similar to our *in vitro* transcription-translation reactions (39 mM HEPES [pH 7.5], 7.7 mM magnesium acetate, 1.3 mM magnesium chloride, 13 mM potassium chloride, 65 mM potassium glutamate, 0.65 mM dithiothreitol (DTT), 3.9 % glycerol). 94 ± 6% of the biotinylated DNA was associated with the beads compared to 3 ± 4% non-specifically associated by non-biotinylated DNA (Fig. S5).

**Fig S5. Streptavidin association is specific for biotinylated DNA.** Biotinylated or non-biotinylated DHFR DNA template was bound to streptavidin beads, followed by separation of the bound and free fractions. The DNA was electrophoresed and the percent of DNA in each fraction was quantified by densitometry.

dCas9 blocks were assembled *in vitro* by incubating 210 nM DNA template encoding 5’ biotinylated DHFR with 1.02 µM dCas9 and 1.02 µM sgRNA at 37°C for 20 min. *In vitro* transcription and translation were initiated by addition of energy solution (at the concentration indicated for *in vitro* transcription), OnePot translation factor mix (1.7 mg/mL), *E. coli* RNA polymerase holoenzyme (15 nM), release factor mix containing RF1, RF2, and RF3 (23 ng/µL), EF-Tu (10 µM), ribosomes (1.4 μM) to yield a final DNA template concentration of 60 nM. Reactions were incubated at 37°C for 2 min, followed by addition of heparin to 100 µg/mL. Reactions were incubated for 18 min and bound to streptavidin beads. Bound complexes were released from the beads by incubating with 30 mM free biotin for 10 min, followed by heating in 4X Laemmli buffer at 95°C for 10 min. RNA polymerase in the bead and supernatant fractions was detected by western blotting with α-RNA polymerase β antibody (Abcam, Cambridge, England; cat. ab191598) and goat anti-rabbit IgG StarBright Blue 700 (Bio-Rad). Blots were visualized in a ChemiDoc MP. We confirmed that the biotinylated template produced a comparable amount of protein to the non-biotinylated template by [^35^S]-methionine incorporation as described above (Fig. S6).

**Fig S6. Biotinylation of DHFR template DNA does not affect translation yield.** DHFR template DNA with or without 5’ biotinylation was used as template for *in vitro* transcription and translation as described in the methods. Translation was monitored by incorporation of [^35^S]-methionine and detected by phosphorimaging. The amount of DHFR protein produced from biotinylated DHFR template was calculated as a percentage relative to non-biotinylated template DNA.

GalR blocks were assembled by incubating 106 nM DHFR containing the GalR operator site with 8.5 µM GalR in energy solution (at the concentrations indicated for *in vitro* transcription) for 20 min. *In vitro* transcription and translation were initiated by addition of OnePot translation factor mix (1.7 mg/mL), *E. coli* RNA polymerase holoenzyme (15 nM), release factor mix containing RF1, RF2, and RF3 (23 ng/µL), EF-Tu (10 µM), and ribosomes (1.4 μM) to yield a final DNA template concentration of 60 nM. Incubation of the translation reactions and separation of bead and supernatant fractions were performed as described for dCas9.

### mRNA release assay

For measurement of transcript release without translation, dCas9 blocks were assembled by incubating 244 nM biotinylated DHFR DNA template with 1.22 µM dCas9 and 1.22 µM sgRNA in dCas9 reaction for 20 min at 37°C. Energy solution (at the concentrations indicated for *in vitro* transcription) and 314 nM *E. coli* RNA polymerase holoenzyme were added to make a final DNA template concentration of 148 nM. Reactions were incubated for 20 min at 37°C, followed by incubation with an equal volume of Dynabeads MyOne Streptavidin C1 magnetic beads (Invitrogen) for 5 min. The bound and supernatant fractions were collected, DNase I treated, heated to 65°C in urea loading buffer, and resolved on 5% denaturing urea-PAGE. Transcripts were visualized by ethidium bromide staining and imaged on a ChemiDoc MP (Bio-Rad).

For measurement of transcript release with translation, dCas9 blocks were assembled by incubating 108 nM 5’ biotinylated DHFR DNA template with 550 nM dCas9 and 550 nM sgRNA at 37°C for 20 min. Reactions were added to energy solution (60 mM HEPES [pH 7.5], 14 mM magnesium acetate, 120 mM potassium glutamate, 1.2 mM dithiothreitol (DTT), 24 mM creatine phosphate, 24 µM folinic acid, 2.4 mM spermidine, 1.1 mM ATP, 1.1 mM GTP, 1.1 mM CTP, 0.05 mM UTP, amino acid mix (0.24 mM methionine and the other 19 common amino acids at 2.4

mM), 3.1 mg/mL tRNA) to make a final DNA template concentration of 50 nM. Transcription was initiated with 100 nM *E. coli* RNA polymerase holoenzyme and transcripts were labeled with [α-^32^P]-UTP (0.24 μCi, 3000 Ci/mmol) (Revvity). After 5 min incubation at 37°C, reactions were bound to an equal volume of Dynabeads MyOne Streptavidin C1 magnetic beads (Invitrogen) for 5 min. The beads were separated from supernatant and resuspended in energy solution without any nucleoside triphosphates. The reactions were split into equal volumes and added to OnePot translation factor mix (2.2 mg/mL), release factor mix containing RF1, RF2, and RF3 (30 ng/µL), EF-Tu (12 µM), ribosomes (1.8 µM), 246 nM non-biotinylated DNA competitor, and either ATP (3.4 mM) and GTP (3.4 mM) or 68 nM HEPES [pH 7.5] as a negative control. Reactions were incubated at 37°C for 10 min, followed by binding to streptavidin beads. Because released transcript was rapidly degraded in the supernatant (Fig. S7), we measured the amount of transcript that was retained on the beads. The bead fractions were collected, and incubated with 95% formamide, 10 mM EDTA at 65°C for 5 min, and the supernatant was analyzed on urea-polyacrylamide gels and visualized by phosphorimaging. Quantification of band intensity was performed in ImageJ.

**Fig. S7. Released transcript is rapidly degraded *in vitro*.** Biotinylated DHFR template DNA without dCas9-sgRNA was transcribed *in vitro* and then incubated with streptavidin beads. The supernatant containing released transcript was split into equal volumes. Translation factors and ribosomes were added to one of the split reactions and an equal volume of transcription buffer was added to the other reaction. Both reactions were incubated at 37C for 10 min, electrophoresed, and phosphorimaged.

### In vivo trans-translation

X90*ssrA*H6D or X90Δ*ssrA*::cat with pTrc99a-2XFLAGEGFP and pBAD-dCas9sgRNA were grown to OD_600_ ∼0.6 and induced with 1 mM IPTG (for 2XFLAGEGFP), 0.5 µg/uL anhydrotetracycline (for dCas9), and/or 13 mM arabinose (for sgRNA) for 3.5 h at 37°C with agitation. Cells were harvested by centrifugation and resuspended in a volume of Laemmli buffer proportional to the wet cell pellet mass. Western blots were probed with either α-FLAG M2 (Thermo Fisher Scientific, cat. F1804, 1:2000 dilution) or α-{Citation} (Qiagen, Hilden, Germany; cat. 34660, 1:3000 dilution), followed by α-mouse IgG HRP conjugate (Bio-Rad, cat. 170-6516, 1:3000 dilution). Blots were visualized in a ChemiDoc MP (Bio-Rad).

### *In vivo trans*-translation in mitomycin treated cells

X90*ssrA*H6D was grown to OD_600_ ∼0.4 at 37°C with agitation. Mitomycin C (Enzo, Farmingdale, NY) was added to 2 µg/mL and the cultures were incubated for 5 min at 37 °C with agitation. Cells were harvested by centrifugation and resuspended in a volume of Laemmli buffer proportional to the wet cell pellet mass. Western blots were probed with α-Penta-His (Qiagen), followed by α-mouse IgG HRP conjugate (Bio-Rad). Quantification of band intensity was performed in ImageJ and the percent tagging with MMC treatment was calculated as the intensity in the +MMC lane divided by the -MMC lane, which was set to a baseline of 100%.

### Mass spectrometry

To determine the identity of the tagged truncated product produced from ribosome-RNA polymerase-dCas9 collision, X90*ssrA*H6D expressing pTrc99a-2XFLAGEGFP and pBAD-dCas9gRNA was induced as above. FLAG-tagged proteins were purified with Anti-DYKDDDDK magnetic agarose (Thermo Fisher Scientific, cat. 36797) according to manufacturer

recommendations. The eluted proteins were digested with Asp-N (Promega, Madison, WI) and analyzed by LC-MS/MS on a Thermo Orbitrap Ascend (University of Texas at Austin Center for Biomedical Research Support Biological Mass Spectrometry Facility (RRID:SCR_021728)). Sample analysis was performed in Sequest (Thermo Fisher Scientific, version IseNode in Proteome Discoverer 2.5.0.400) with a fragment ion mass tolerance of 0.60 Da and a parent ion tolerance of 10 PPM. The peptide fragments were searched against a list of all possible sequences for translation of additional codons past the expected ribosomal P-site, each of which would produce a unique peptide. Peptide identification was performed in Scaffold (Proteome Software Inc., Portland, OR; version Scaffold_5.3.4) with a peptide identification criteria of >80% probability by Percolator posterior error probability calculation (63), and a protein identification criteria of >12% probability, <1.0% false discovery rate, and detection of at least one identified peptide. Protein probabilities were determined by the Protein Prophet algorithm (64). Peptides that could have been generated from more than one protein were assigned based on the principles of parsimony.

## Acknowledgements

We thank Jason Peters for sharing pJMP1337 and for helpful discussions, Christopher Hayes for the gift of X90*ssrA*H6D, Neeraja Marathe for constructing the glyQS and pheST plasmids, and Audrey Long for constructing X90Δ*ssrA*::*cat*.

