## Supplemental figures for "Blocked transcription-translation complexes are rescued by transcript release followed by *trans*-translation"

Table S1.

| Peptide sequence | Sequest Xcorr[a] |
|---|---|
| DDGNYKTRAEVKFEG | 2.53 |
| DFFKSAMPEGYVQERTIFFK | 2.74 |
| DFFKSAMPEGYVQERTIFFKD | 3.12 |
| DFFKSAMPEGYVQERTIFFKDDGNYKTRAEVKFEG | 1.67 |
| DGNILGHKLEYNYNSHNVYIMA | 3.77 |
| DGNYKTRAEVKFEG | 2.62 |
| DGPVLLPDNHYLSTQSALSK | 3.89 |
| DGPVLLPDNHYLSTQSALSKDPNEKR | 3.7 |
| DHYQQNTPIGDGPVLLP | 4.73 |
| DHYQQNTPIGDGPVLLPDNHYLSTQSALSK | 4.68 |
| DKQKNGIKVNFKIRHNIE | 2.04 |
| DNHYLSTQSALSK | 3.55 |
| DNHYLSTQSALSKDPNEKR | 3.72 |
| DTLVNRIELKGIDFKE | 3.72 |
| DTLVNRIELKGIDFKEDGNILGHKLEYNYNSHNVYIMA | 1.23 |
| DDGNYKTRAEVKFEG | 2.53 |
| DFFKSAMPEGYVQERTIFFK | 2.74 |
| DFFKSAMPEGYVQERTIFFKD | 3.12 |
| DFFKSAMPEGYVQERTIFFKDDGNYKTRAEVKFEG | 1.67 |
| DGNILGHKLEYNYNSHNVYIMA | 3.77 |
| DGNYKTRAEVKFEG | 2.62 |
| DKQKNGIAANDHHHHHHD | 1.72 |
| DTLVNRIELKGIDFKE | 3.72 |
| DTLVNRIELKGIDFKEDGNILGHKLEYNYNSHNVYIMA | 1.23 |

[a]The cross-correlation score between acquired and theoretical spectra

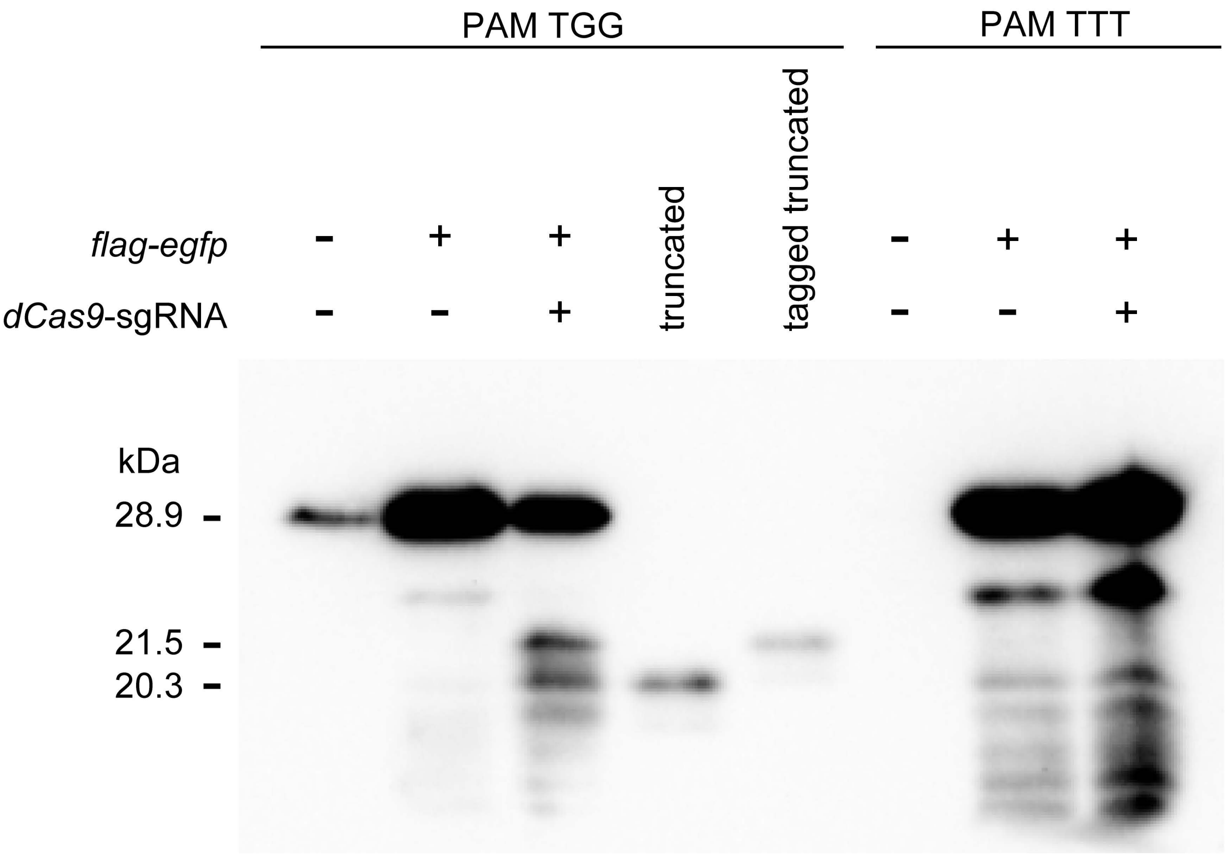
PAM TGG
PAM TTT
flag-egfp
dCas9-sgRNA
truncated
tagged truncated
kDa
28.9
21.5
20.3

Figure S1.

| B | B Ions | B+2H | B-NH3 | B-H2O | AA | Y Ions | Y+2H | Y-NH3 | Y-H2O | Y |
|---|---|---|---|---|---|---|---|---|---|---|
| 1 | 116.0 | 58.5 | | 98.0 | D | 2,181.2 | 1,091.1 | 2,164.2 | 2,163.2 | 18 |
| 2 | 244.1 | 122.6 | 227.1 | 226.1 | K | 2,066.2 | 1,033.6 | 2,049.2 | 2,048.2 | 17 |
| 3 | 372.2 | 186.6 | 355.2 | 354.2 | Q | 1,938.1 | 969.6 | 1,921.1 | 1,920.1 | 16 |
| 4 | 500.3 | 250.6 | 483.3 | 482.3 | K | 1,810.0 | 905.5 | 1,793.0 | 1,792.0 | 15 |
| 5 | 614.3 | 307.7 | 597.3 | 596.3 | N | 1,681.9 | 841.5 | 1,664.9 | 1,663.9 | 14 |
| 6 | 671.3 | 336.2 | 654.3 | 653.3 | G | 1,567.9 | 784.5 | 1,550.9 | 1,549.9 | 13 |
| 7 | 784.4 | 392.7 | 767.4 | 766.4 | I | 1,510.9 | 755.9 | 1,493.9 | 1,492.9 | 12 |
| 8 | 912.5 | 456.8 | 895.5 | 894.5 | K | 1,397.8 | 699.4 | 1,380.8 | 1,379.8 | 11 |
| 9 | 1,011.6 | 506.3 | 994.6 | 993.6 | V | 1,269.7 | 635.4 | 1,252.7 | 1,251.7 | 10 |
| 10 | 1,125.6 | 563.3 | 1,108.6 | 1,107.6 | N | 1,170.6 | 585.8 | 1,153.6 | 1,152.6 | 9 |
| 11 | 1,272.7 | 636.9 | 1,255.7 | 1,254.7 | F | 1,056.6 | 528.8 | 1,039.6 | 1,038.6 | 8 |
| 12 | 1,400.8 | 700.9 | 1,383.8 | 1,382.8 | K | 909.5 | 455.3 | 892.5 | 891.5 | 7 |
| 13 | 1,513.9 | 757.4 | 1,496.9 | 1,495.9 | I | 781.4 | 391.2 | 764.4 | 763.4 | 6 |
| 14 | 1,670.0 | 835.5 | 1,653.0 | 1,652.0 | R | 668.3 | 334.7 | 651.3 | 650.3 | 5 |
| 15 | 1,807.0 | 904.0 | 1,790.0 | 1,789.0 | H | 512.2 | 256.6 | 495.2 | 494.2 | 4 |
| 16 | 1,921.1 | 961.0 | 1,904.1 | 1,903.1 | N | 375.2 | 188.1 | 358.2 | 357.2 | 3 |
| 17 | 2,034.2 | 1,017.6 | 2,017.1 | 2,016.2 | I | 261.1 | 131.1 | | 243.1 | 2 |
| 18 | 2,181.2 | 1,091.1 | 2,164.2 | 2,163.2 | E | 148.1 | 74.5 | | 130.0 | 1 |

Figure S2.

| B | B Ions | B+2H | B-NH3 | B-H2O | AA | Y Ions | Y+2H | Y-NH3 | Y-H2O | Y |
|---|---|---|---|---|---|---|---|---|---|---|
| 1 | 116.0 | 58.5 | | 98.0 | D | 2,111.0 | 1,056.0 | 2,093.9 | 2,093.0 | 18 |
| 2 | 244.1 | 122.6 | 227.1 | 226.1 | K | 1,995.9 | 998.5 | 1,978.9 | 1,977.9 | 17 |
| 3 | 372.2 | 186.6 | 355.2 | 354.2 | Q | 1,867.8 | 934.4 | 1,850.8 | 1,849.8 | 16 |
| 4 | 500.3 | 250.6 | 483.3 | 482.3 | K | 1,739.8 | 870.4 | 1,722.8 | 1,721.8 | 15 |
| 5 | 614.3 | 307.7 | 597.3 | 596.3 | N | 1,611.7 | 806.3 | 1,594.7 | 1,593.7 | 14 |
| 6 | 671.3 | 336.2 | 654.3 | 653.3 | G | 1,497.6 | 749.3 | 1,480.6 | 1,479.6 | 13 |
| 7 | 784.4 | 392.7 | 767.4 | 766.4 | I | 1,440.6 | 720.8 | 1,423.6 | 1,422.6 | 12 |
| 8 | 855.5 | 428.2 | 838.4 | 837.5 | A | 1,327.5 | 664.3 | 1,310.5 | 1,309.5 | 11 |
| 9 | 926.5 | 463.8 | 909.5 | 908.5 | A | 1,256.5 | 628.8 | 1,239.5 | 1,238.5 | 10 |
| 10 | 1,040.5 | 520.8 | 1,023.5 | 1,022.5 | N | 1,185.5 | 593.2 | 1,168.4 | 1,167.5 | 9 |
| 11 | 1,155.6 | 578.3 | 1,138.5 | 1,137.6 | D | 1,071.4 | 536.2 | | 1,053.4 | 8 |
| 12 | 1,292.6 | 646.8 | 1,275.6 | 1,274.6 | H | 956.4 | 478.7 | | 938.4 | 7 |
| 13 | 1,429.7 | 715.4 | 1,412.7 | 1,411.7 | H | 819.3 | 410.2 | | 801.3 | 6 |
| 14 | 1,566.8 | 783.9 | 1,549.7 | 1,548.7 | H | 682.3 | 341.6 | | 664.3 | 5 |
| 15 | 1,703.8 | 852.4 | 1,686.8 | 1,685.8 | H | 545.2 | 273.1 | | 527.2 | 4 |
| 16 | 1,840.9 | 920.9 | 1,823.8 | 1,822.9 | H | 408.2 | 204.6 | | 390.2 | 3 |
| 17 | 1,977.9 | 989.5 | 1,960.9 | 1,959.9 | H | 271.1 | 136.1 | | 253.1 | 2 |
| 18 | 2,111.0 | 1,056.0 | 2,093.9 | 2,093.0 | D | 134.0 | 67.5 | | 116.0 | 1 |

Figure S3.

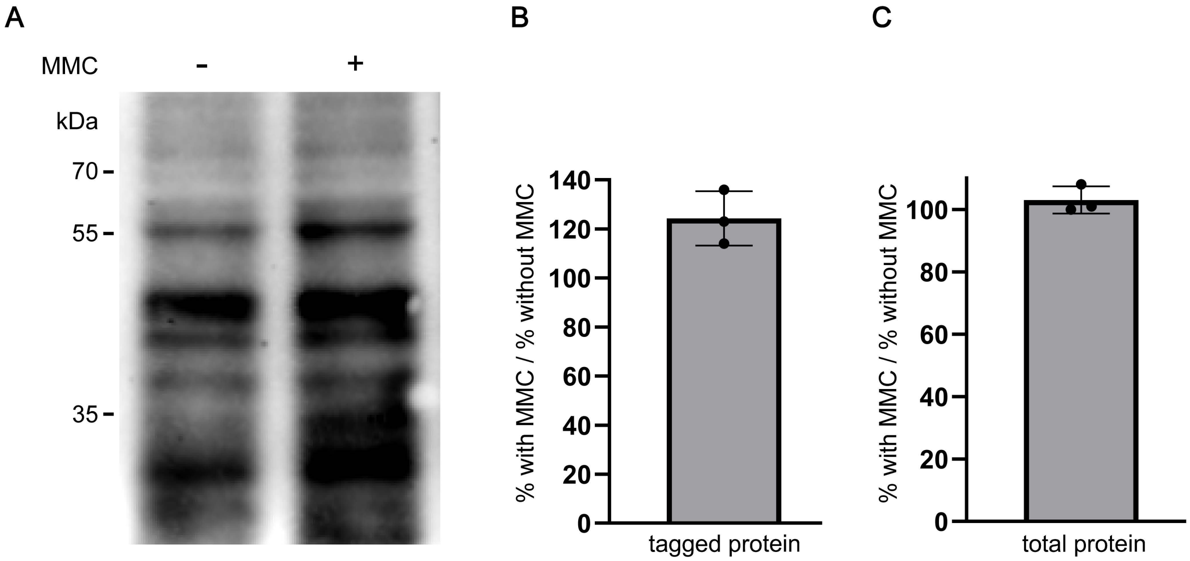
A
MMC
-
+
kDa
70
55
35
B
140
120
100
80
60
40
20
0
% with MMC / % without MMC
tagged protein
C
100
80
60
40
20
0
% with MMC / % without MMC
total protein

Figure S4.

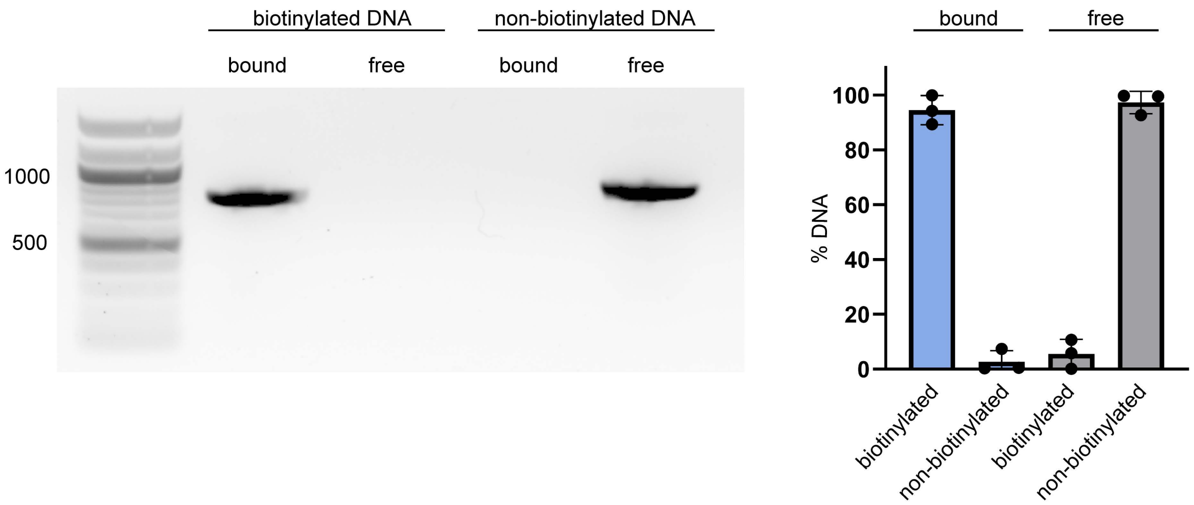

biotinylated DNA
non-biotinylated DNA
bound
free
bound
free
1000
500
bound
free
100
80
60
40
20
0
% DNA
biotinylated
non-biotinylated
biotinylated
non-biotinylated


Figure S5.

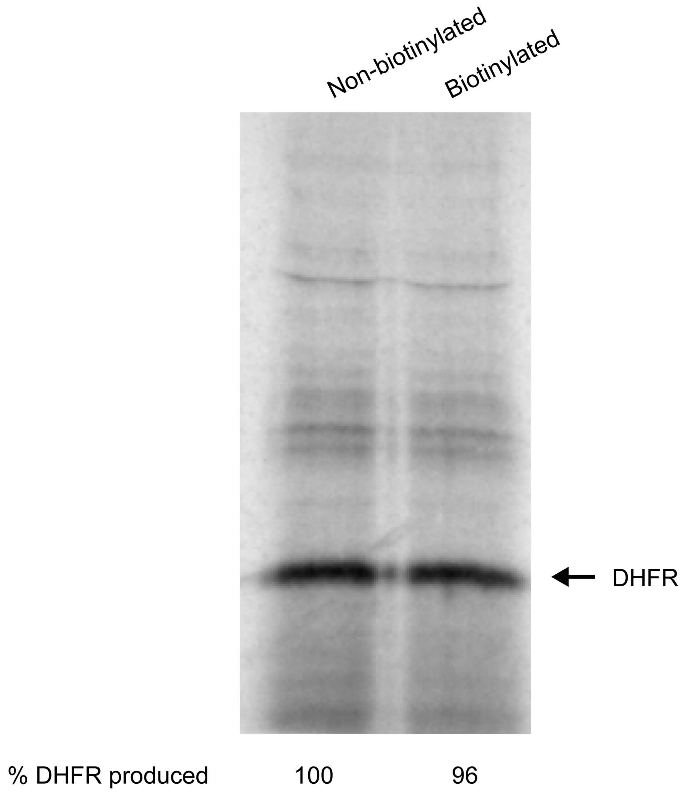


Figure S6.

translation factors - +

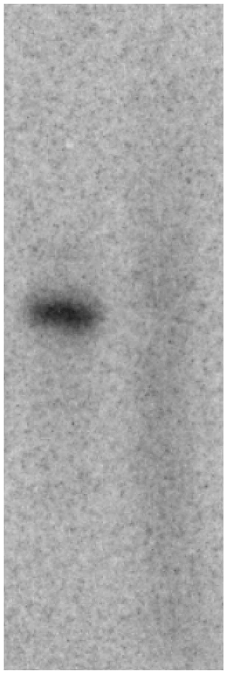

Figure S7.
